# How strong is enemy release? A systematic compilation across taxa and approaches

**DOI:** 10.1101/2023.11.24.568498

**Authors:** Karen Zeng, Jess Schembri, Eve Slavich, Angela T. Moles

**Affiliations:** Evolution & Ecology Research Centre, School of Biological, Earth and Environmental Sciences, UNSW Sydney, NSW 2052, Australia; School of Mathematics and Statistics, UNSW Sydney, Sydney, New South Wales, Australia

**Keywords:** enemy release, biogeographic, community, non-indigenous species, invasive alien species, enemy escape, natural enemies

## Abstract

The enemy release hypothesis posits that introduced species escape some of their predators, pathogens and parasites when they move to a new range. We used a systematic review to compile data from 691 quantifications of enemy release spanning plants, animals and algae in aquatic and terrestrial systems worldwide. Data from 311 biogeographic contrasts (between home and new range) revealed that on average, a species experience only 30% as much enemy pressure in their introduced range as they experience in their native range. In contrast, data from 380 community contrasts (between native and introduced species) revealed that introduced species experience on average 57% of the enemy pressure that their native congeners endure. Interestingly, one third (36%) of contrasts showed higher, rather than lower, enemy pressure on the introduced population. Enemy release was consistently strong in contrasts of the diversity of enemies, intermediate in contrasts comparing enemy damage, and not significant in contrasts of host fitness, suggesting that while introduced populations are attacked by fewer enemies, this does not always result in higher fitness. We also found that enemy release was higher in molluscs and fish but lower in insects and algae, indicating that certain taxa may be favoured by enemy release. We hope that an improved understanding of the extent to which introduced species are released from enemy pressures will help managers to identify good opportunities for biocontrol, and to understand the factors likely to be affecting the success of invasive species.

## Introduction

We are taught as scientists that species become well-adapted to their native environments through the evolutionary pressures of natural selection (Darwin 1864). Yet, every so often we find that recently introduced organisms outperform native species to the point of environmental damage. The success of these species seemingly contradicts our basic understanding of the nature of evolution, and is one of the great conundrums of invasion ecology (Elton 1972). One commonly cited theory that attempts to explain the success of introduced species is the enemy release hypothesis (Keane and Crawley 2002). Also known as the ‘enemy escape’ (Mlynarek et al. 2017) or ‘natural enemies hypothesis’ (Liu and Stiling 2006), the enemy release hypothesis suggests that introduced species often experience a reduction in top-down biotic pressure. This can contribute to some introduced species becoming invasive or otherwise overabundant. This idea that introduced species succeed partly through a lack of enemy pressure also underpins the success of classical biological control, in which specialised enemies are introduced to counterbalance enemy release and reduce the fitness of introduced species (Keane and Crawley 2002). Despite the conceptual and applied importance of enemy release, and the rapid accumulation of information (Bornmann et al. 2021), the enemy release literature as a whole has not been reviewed since 2012 (Heger and Jeschke 2014). Our primary aim is thus to provide a quantitative synthesis of support for the enemy release.

First, we aimed to quantify the average magnitude of enemy release across plant and animal species worldwide, and to determine what proportion of species show significant enemy release. Previous syntheses suggest that enemy release is supported in just over half the studies in which it is measured (Colautti et al. 2004, Heger and Jeschke 2014). However, previous attempts to quantify support for enemy release in literature syntheses have mainly used vote-counting methods. In cases where effect sizes have been used, the scope of studies considered has been limited to individual host-enemy systems such as plant-fungal/viral pathogen, plant-herbivore and animal-parasite (Mitchell and Power 2003, Torchin et al. 2003, Liu and Stiling 2006, Meijer et al. 2016). Thus, we aim to perform the first broad, quantitative assessment of support for the enemy release hypothesis.

Studies of enemy release usually use one of two major methodological approaches, which have been termed biogeographic and community enemy release ((Colautti et al. 2004); Fig. 1). Studies of **biogeographic enemy** *release* compare the amount of enemy pressure a species experiences in its introduced range to the amount of enemy pressure the same species experiences in its native range. Studies of **community enemy release** compare enemy pressure to a species in its introduced range with enemy pressure on a nearby native species that is as similar to the host as possible in both taxonomy and traits (i.e. a native congener/confamillial). Interestingly, while the few studies that use both methods do not consistently find that one produces stronger support for enemy release than the other (Norghauer et al. 2011, Meijer et al. 2016, Berthelot et al. 2023), reviews of enemy release literature find that biogeographic studies support enemy release more often than do community studies (Colautti et al. 2004, Heger and Jeschke 2014). Despite the considerable effect that experimental approach may have on results, individual studies often ignore the distinction when drawing conclusions about invasive potential (Stutz et al. 2016, Najberek et al. 2020a), or acknowledge the difference while being unable to address it (Davidson et al. 2023). We thus quantified the magnitude of enemy release when measured with both community and biogeographic approaches.

**Figure 1.**
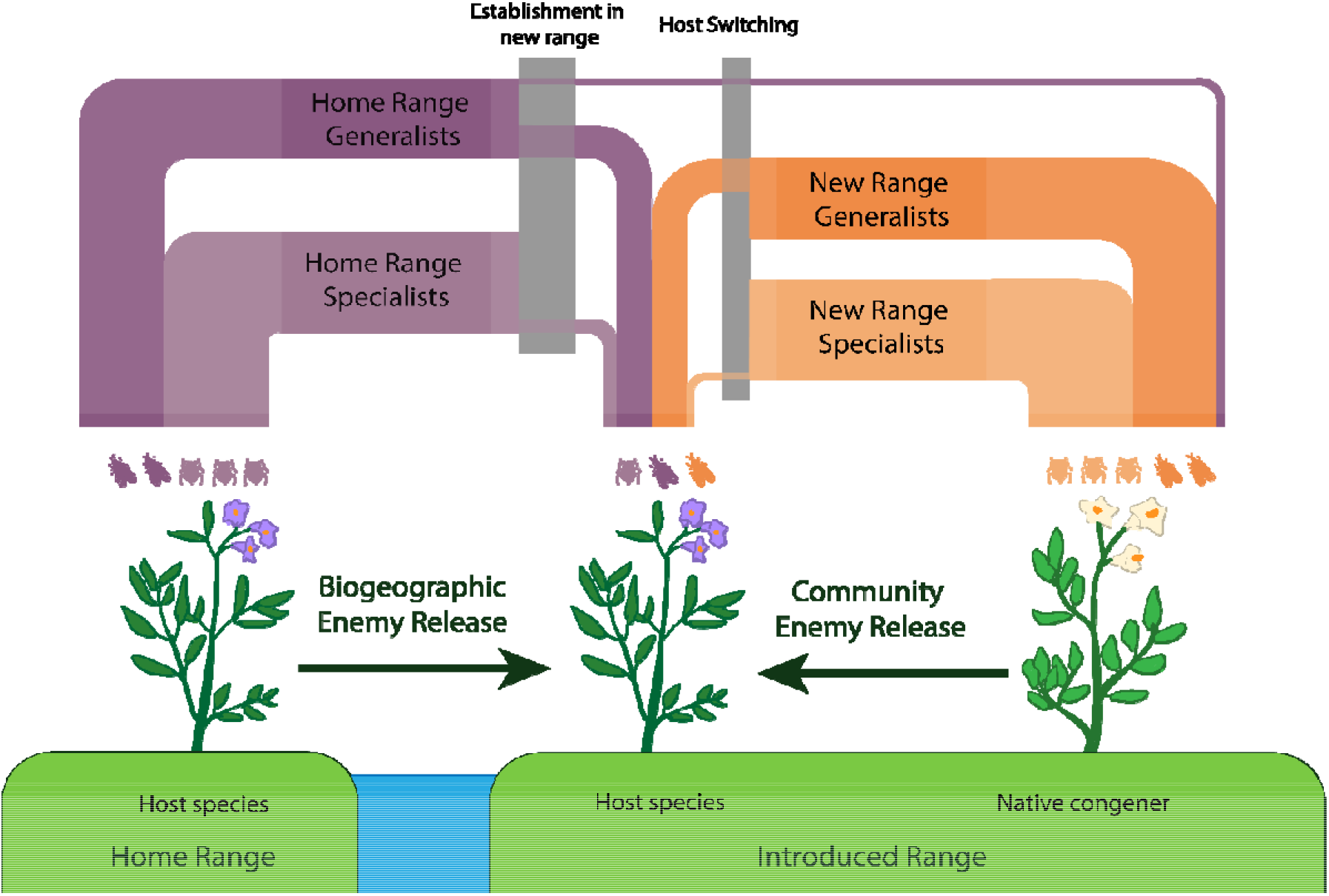
The two main ways to measure enemy release are the biogeographic and community methods. Introduced species may be released from enemy pressure in their new range either by leaving behind old enemies, or by avoiding new range enemies which many switch from attacking native hosts to the new introduced species.

Our second goal was to determine whether the degree of enemy release observed varies between studies that use different measures of enemy pressure. Some studies measure enemy pressure by quantifying biomass loss or by looking at the number and diversity of enemies predating on individuals (Keane and Crawley 2002, Lester et al. 2015, Najberek et al. 2020b). Other studies measure host fitness via total host biomass (Fan et al. 2013), offspring survival or offspring/seed weight (Yan and Chen 2007). Finally, some studies look for changes in species’ defence traits (Wan et al. 2019) as predicted under the shifting defence hypothesis (Müller-Schärer et al. 2004), or trade-offs between defence/tolerance and competition (Franks et al. 2008, Li et al. 2012) as predicted by the evolution of increased competitive ability theory (Blossey and Notzold 1995, Joshi and Vrieling 2005). Differences in support between these measurements will reveal to what extent a decreased diversity of enemies flows on to decreased damage and higher fitness. Previously syntheses have found that enemy release studies measuring damage more often support the predictions of the enemy release hypothesis than do studies measuring host performance (Heger and Jeschke 2014) but have yet to compare the magnitude of support between measurements. We predict that enemy diversity would produce the most evidence for enemy release and that enemy damage, host fitness and evolutionary change would subsequently show less support for the hypothesis as they become less directly affected by the barriers of establishment and host-switching that cause enemy release.

Next, we asked whether the magnitude of enemy release varied with the taxonomic group of the introduced species. While enemy release is a universal concept that can be applied to any species range shift scenario, the study of enemy release has maintained close connections to its roots of terrestrial plant-herbivore interactions. Heger and Jeschke (2014) found that the proportion of studies that supported the enemy release hypothesis substantially differed between taxa; and that studies of plants made up approximately 80% of the literature. Variability between study systems have been used to explain contradictions between studies has been used as a reason why cross-taxa studies are avoided (Meijer et al. 2016), despite strong calls for such studies (Pyšek et al. 2008). Indeed, the variation in support for enemy release between taxa may help inform us of what types of introduced species are more likely to benefit from enemy release.

Finally, we test the hypothesis that enemy release is stronger in marine than in terrestrial habitats. This prediction is based on the observation that herbivores consume three to four times the amount of living plant in marine systems than on land, indicating that enemies play a larger role in controlling biomass in marine environments (Shurin et al. 2005), and therefore that release from such enemies may be additionally beneficial to introduced marine species. On the other hand, aquatic systems contain more generalist predators/herbivores (Shurin et al. 2005), and diseases and predators can spread more freely in marine ecosystems (McCallum et al. 2004) which may reduce the importance of enemy release in underwater systems. To our knowledge, there has yet to be a comparison of enemy release between terrestrial and aquatic systems.

By quantifying broad patterns in the strength of enemy release between taxa and environments, we hope to identify generalisations that will improve understanding of biological invasions and facilitate more effective biocontrol efforts.

## Methods

We performed a systematic literature search for papers with the term “enemy release” or the terms “enemy” and “release” on Web of Science and Scopus up to May 2020 (exact search string available in Appendix 1).

We included any study which quantified the enemy pressure in at least one introduced population of a specified species, and at least one control group. In biogeographic contrasts, the control group is a population of the same species in its home range. In community contrasts, the control group is a native species that is as phylogenetically and functionally similar to the introduced species as possible. We randomised the order of papers and used only titles, keywords and abstracts in manual screening to determine whether paper was likely to be relevant. We excluded biocontrol efficacy reports, as while classical biological control is often used as an example for enemy release, the action of choosing the most appropriate biocontrol agent would naturally bias the results towards a favourable outcome, thereby making the success of biological control a poor measure for quantifying the enemy release on any particular host (Keane and Crawley 2002). We also decided to exclude enemy release records observed exclusively in agricultural systems because the conditions present including the enforcement of monocultures and lack of resource limitation may not be applicable to natural ecosystems. Meta analyses and reviews were excluded but noted down in the manual screening process for reference. For PRISMA reporting documents and additional methods see supplementary file 3.

Full papers were automatically downloaded, and missing papers were manually downloaded if available online. KZ and JS manually read each paper and recorded measurements of enemy release. Papers within the final dataset were checked for retractions, addendums and errata using the Retraction Watch Database (http://retractiondatabase.org/). No retraction issues were identified, but we did identify ‘publication overlap’ (Urbanowicz and Reinke 2018) where multiple publications used the same dataset that resulted in the same data being counted three times, which we rectified by excluding all but one copy.

From each paper, we obtained a measurement of the strength of enemy pressure affecting each measured population.

Enemy release was quantified as a log response ratio between enemy pressure experienced by an introduced population and its control group. We also reported back-transformed averages in the results to better communicate the scale of enemy release observed.

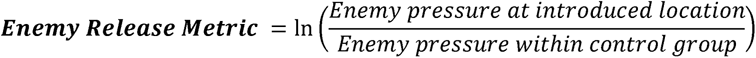

This response ratio allows for enemy release to be quantified intuitively with positive numbers indicating higher enemy release. For example, if a host organism experiences half as much damage in their introduced range than in their native range we would allocate them a positive enemy release value of 0.7. If a host organism experiences no enemy release (i.e. enemy pressure is the same in the introduced population as in the control group; 1/1) it would have an enemy release metric of 0. Finally, if there is greater enemy pressure in the new range it would have a negative enemy release metric.

We recorded individual enemy release measurements for each introduced population of all species for which sufficient data were available in the literature. If a study provided measurements for multiple introduced populations, we created a new record for each introduced population. If a study sampled from multiple control populations, we took the average of each. If there were multiple measurements of the same enemy-host interaction we would prioritise the one with the most direct fitness consequence to the introduced species (i.e., seed damage > total biomass change).

We recorded additional information from the literature including study species, environment and methodological choices. We categorised introduced species into taxonomic groups using additional information obtained from the NCBI Taxonomy Database (https://www.ncbi.nlm.nih.gov/taxonomy). We assumed that coordinates were in WGS84 unless otherwise specified, in which case they were converted to WGS84. In cases where locations were given in formats other than coordinates, we used Google Earth to manually determine central point of a locations (i.e., the midpoint of a named creek or the centroid of a named town). In cases where location was very uncertain (radius of > 50km) we excluded the record from our data. Complete information on inclusion requirements is available in supplementary file 2 with a reference list for included publications in supplementary file 3.

We obtained enemy release data for 691 population*species combinations, from 223 published data sources (available in supplementary file 1). Of these sources, a quarter of studies measured enemy release incidentally, with the rest including some test of enemy release in their main hypothesis (155/217). Interestingly, only 5 out of 217 studies included in our dataset measured both biogeographic and community enemy release.

## Data analysis

We first tested whether to consider biogeographic and community enemy release as separate metrics by checking if the strength of their log response ratio contrasts differed using a student’s t test. Because we found a substantial significant difference between the two approaches, we performed all subsequent analyses separately for community vs biogeographic enemy release.

To analyse whether enemy release differed based on the type of response variable used, we performed an ANOVA between the strength of enemy release and whether a study looked at enemy diversity, enemy pressure, reductions in host fitness or evolutionary changes in the host. Where evidence for differences between groups was found (p < 0.05), we followed up with a post-hoc Tukey test to identify which groups were driving the effect.

We then tested whether enemy release differed based on the type of host measured. We first produced a summary of sample sizes in order to help determine how to group host organisms. Whenever a group consisted of fewer than three unique studies, we tried to incorporate it into a larger group but otherwise dropped it from further analysis. Groups were chosen based on general morphology and role in the ecosystem rather than taxonomy sensu stricto. For example, birds were separated from other sauropsids due to morphology and algae were included with the plants in the higher level group for photosynthetic ability. To analyse the effect of host on enemy release strength, we performed an ANOVA. As before, we performed a post-hoc Tukey test to determine which groups were different when we found significant differences between groups.

Finally, we tested whether enemy release is stronger in marine environments than in terrestrial or freshwater by performing an ANOVA, calculating summary statistics, and performing a Tukey test if significant differences were found.

We filtered literature and performed analyses using R version 4.2.1 (R Core Team, 2022) and the following additional packages. Papers were downloaded automatically using ‘fulltext’ (Chamberlain 2019), deduplication and abstract screening performed using ‘revtools’ (Westgate 2019) and graphics produced using ‘ggplot2’(Wickham 2016). Code used in this study is available on GitHub (https://github.com/karenZENG-eco/Enemy-Release-Literature-Compilation).

## Results

We found evidence for a difference in the strength of enemy release between records collected using biogeographic contrasts and those collected using community contrasts (t(686) = -4.261, p < 0.0001). The 311 biogeographic contrasts revealed that populations of species in their introduced range experience just 43% of the enemy pressure that populations of the same species endure in their home range (effect size = 0.838, SD = 1.55). In contrast, the 380 community contrasts (between an introduced population and nearby counterparts) indicate that introduced species experience 70% of the enemy pressure experienced by native species (effect size = 0.367, SD = 1.35). Because of this evidence for a substantial difference between the two study types, we performed all subsequent analyses separately for community vs biogeographic contrasts.

### Biogeographic Enemy Release

The metric used to quantify enemy pressure affected the amount of enemy release found in biogeographic contrasts (*F*(3,376) = 5.90, *p* < 0.001). Measures of enemy diversity (effect size = 0.79 ± SD = 1.58, N = 85), enemy damage (effect size = 0.96 ± 1.50, N = 200) and host evolutionary change (effect size = 1.19 ± 1.93, N = 51) suggested that introduced populations experienced around half of the enemy pressure experienced by populations in their native range. Surprisingly, contrasts of host fitness between populations in their introduced and native range found less than 1 percent difference in enemy pressure (effect size = -0.008 ± 0.71, N = 44). Post-hoc Tukey tests confirmed that host fitness differed from all other groups, which did not significantly differ between each other otherwise (Fig. 2a).

**Figure 2:**
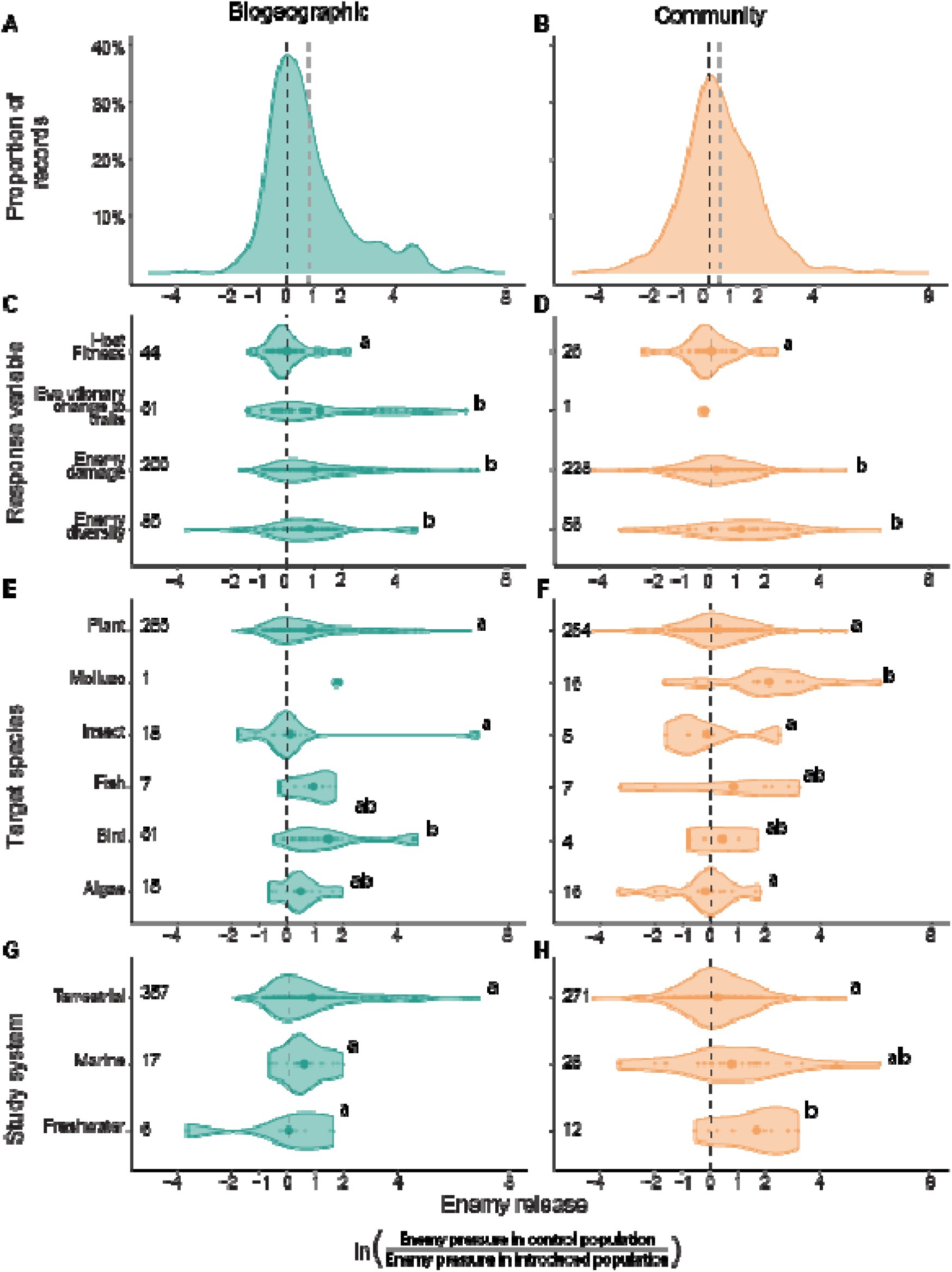
Violin plots showing the distribution of biogeographic and community enemy release. Hollow dots represent individual records, larger solid dots represent group mean, numbers on the left indicate sample size and letters indicate significantly different groups.

The strength of biogeographic enemy release differed between taxonomic groups (*F*(4,371) = 3.59, *p* = 0.007). Birds experienced the greatest reduction in enemy pressure in their introduced range (23% of their native range pressure, effect size = 1.48 ± 1.52, N = 51) while insects barely experienced any enemy release (effect size = 0.11 ± 1.82, N = 18). On average, populations of introduced fish experienced 40% of the enemy pressure experienced by populations of the same species in their home range (effect size = 0.92 ± 0.73, N = 7), while introduced plants experienced 56% (effect size = 0.81 ± 1.52, N = 285), and introduced algae 37% (effect size = 0.46 ± 0.75, N = 15).

Counter to our final prediction, the strength of biogeographic enemy release did not differ between terrestrial (effect size = 0.86 ± 0.47, N = 357), marine (effect size = 0.59 ± 0.44, N = 17) or freshwater (effect size = 0.03 ± 1.93, N = 6) environments (F(2,377) = 1.10 , p = 0.355)) (Fig. 2g).

### Community Enemy Release

The strength of community enemy release differed depending on whether a record measured damage on the host, the fitness of the host, the diversity of the enemies or evolutionary changes in the host (*F*(3,307) = 7.49, *p* < 0.0001)). This result is consistent with results for biogeographic records (Fig. 2). Community comparisons of enemy diversity revealed that introduced species are attacked by only a third of the enemy diversity experienced by their native counterparts (effect size = 1.11, *SD* = 1.79). Unlike what we found in biogeographic enemy release, contrasts that measured enemy damage (effect size = 0.22, *SD* = 1.21) or fitness (effect size = 0.02, *SD* = 0.91) revealed much weaker enemy release (Fig. 2f). We had only a single datapoint for evolutionary change in traits in community contrasts, so did not include it in the above analysis.

The strength of community enemy release differed between taxa (*F*(5,295) = 7.11, *p* < 0.0001)). Molluscs showed the highest average enemy release compared to native congeners, experiencing only 12% of the enemy pressure of native counterparts(effect size = 2.11 ± 1.89, N = 16). The next strongest effects were for fish (44%, effect size = 0.82 ± 2.49, N = 7), birds (65%, effect size = 0.42 ± 1.13, N = 4), plants (78%, effect size = 0.25 ± 1.20, N = 254) with insects (113%, effect size = -0.13 ± 1.58, N = 5) and algae (120%, effect size = -0.19 ± 1.32, N = 15) experiencing slightly greater enemy pressure than their counterparts (Fig 2g).

In contrast to biogeographic contrasts, community contrasts of enemy release revealed evidence for a difference between terrestrial, marine and freshwater studies (*F*(2,308) = 7.869, *p* < 0.001)). Freshwater ecosystems had substantially stronger enemy release (effect size = 1.67 ± 1.28, N = 12) than either marine (effect size = 0.77 ± 2.14, N = 28) or terrestrial systems (effect size = 0.26 ± 1.21, N = 271).

## Discussion

The strength of enemy release differs substantially based on how the enemy release hypothesis is tested. On average, an introduced population experiences 43% as much enemy pressure as a population of the same species in its native range (biogeographic contrast), and 70% as much pressure as matched native species in the invaded range (community contrasts). We can think of two non-exclusive reasons for why the biogeographic contrasts show almost twice the effect size of the community contrasts. Firstly, the relative release from enemies between old and introduced ranges may be more important to initial establishment than the relative release from enemies within the new range community, which may instead be more important for long term performance. Thus, it may be that unless a species is by chance very well pre-adapted to the environment to which it is introduced, there may be some threshold level of biogeographic enemy release required for a species to establish. Introductions which fail to establish cannot be sampled, and so we would find that biogeographic enemy release is higher whereas the slower-acting influence of community enemy release remains unchanged. Secondly, biogeographic contrasts compare within the same species across different environments, whereas community contrasts hold the environment constant, but compare species that differ in many different traits (Meijer *et al*., 2015). If native and successfully introduced species consistently differ in traits (Van Kleunen *et al*., 2010; Montesinos, 2022) and if some of these traits affect susceptibility to enemy pressure (such as fast growth strategies), then this would lower the average magnitude of enemy release observed in community, but not biogeographic studies.

In both biogeographic and community studies the strongest evidence for enemy release was found in contrasts that measured enemy diversity. This lower diversity of enemies on introduced populations is consistent with a loss of enemies as predicted under the enemy release hypothesis (Fig. 1) (Keane & Crawley, 2002). The idea that introduced populations shed enemies underpin both the evolution of increased competitive ability theory and the shifting defence hypothesis (Blossey & Notzold, 1995; Joshi & Vrieling, 2005). A reduction in enemy diversity coincided with less enemy damage in biogeographic contrasts (Fig. 2a) but had relatively little effect on enemy damage in community contrasts (Fig. 2b). The drop in the magnitude of enemy release between diversity and damage suggests that the few taxa of enemies that remain may also be the most effective or numerically dominant. One possibility is that the reduction in specialist enemies associated with introduction to a new range (Fig. 1) may be less important to the fitness of introduced species than initially believed (Keane & Crawley, 2002). Existing attempts to differentiate between generalist and specialist effects seem to indicate that generalist enemies can at least equal specialists in their importance, with both *Silene latifolia* (Wolfe, 2002) *Brachypodium sylvaticum* (Halbritter *et al.,* 2017) being denied complete enemy release in the USA by generalist herbivores and pathogens. However, methodological difficulties were stressed in both studies (comparing between feeding guilds, determining cause and effect in observations and defining the degree of specialisation) which preclude a definitive comparison between generalists and specialists, indicating that this area is a knowledge gap well-suited for future studies.

Enemy release was evident, at least in biogeographic studies, when enemy pressure was measured as diversity, damage or trait evolution but not when researchers measured the fitness of the introduced species (Fig. 2c, d). An interesting implication of this finding is that enemy tolerance and plasticity may play a greater role in moderating the effects of enemy release than expected (Li *et al*., 2012). Support for this idea can be found in common garden comparisons between invasive and non-invasive populations which often report greater plasticity in ‘more successful’ invasive populations (Bhattarai *et al*., 2017; Castillo *et al*., 2021).

There were considerable and relatively consistent taxonomic differences in both biogeographic and community enemy release (Fig. 2e,f). An interesting direction for future research would be to identify which strategies and trait combinations are favoured under enemy release, and if the characteristics that allow species to benefit from enemy loss during introduction are also useful in delaying the gain of new enemies. Given that the historical bias towards terrestrial plants has continued into the current literature (see relative sample sizes in Fig. 2), we thus echo calls to resolve the lack of cross-taxonomic studies within the enemy release literature and invasion biology in general (Jeschke *et al*., 2012). Without controlled comparisons between a varied group of introduced species, it may however be possible for overrepresented taxa to dominate the enemy release literature and cause us to draw incorrect conclusions on the phenomena of enemy release as a whole.

## Conclusion

In our study, we found differences in the magnitude of enemy release between different approaches, taxa and measures of enemy pressure which are important to consider when interpreting or comparing the results of studies that do not share methods. Rather than concluding that any method is ‘better’ at identifying enemy release however, contradictory results between biogeographic vs community enemy release, outlying host fitness and trends between taxa can inform our understanding of how introduction into a new range allows species to survive, establish and potentially invade. We hope that these findings inspire further research into patterns and processes in enemy release so that we can better understand species introductions, produce a more unified understanding of invasion biology, and ideally inform more effective management of both introduced and native species.

## Supporting information

Supplementary File 1 Data for study

Supplementary File 2 Readme for data

## Acknowledgements

We acknowledge Z. Xirocostas and G. Chiarenza for additional insights into enemy release from a fieldwork-based perspective. Work was funded by an Australian Government Research Training Program scholarship to K. Z and an Australian Research Council grant (DP190100243) to A.T.M. The authors have declared that no competing interests exist.

## Supplementary Information

**Supplementary File 1:** Enemy release literature synthesis dataset (in separate .csv)

**Supplementary File 2:** README for enemy release literature synthesis dataset (in separate .txt)

**Supplementary File 3:** Methods and additional graphs including search string, PRISMA documentation and references for enemy release dataset.

