## Supplementary File 2 Readme for data for "How strong is enemy release? A systematic compilation across taxa and approaches"

---------------------------------------
I. About the dataset
---------------------------------------
This is the README for the enemy_release_occurence_dataset
for the Enemy Release Literature Compilation Project.

Please contact Karen ( or 0421773446) 
for more details.

----------------------------------------------------
II. List of Papers to Read (combined_biblio_filter3)
----------------------------------------------------
This table is a list of sources of information and a set of rules to determine 
whether we include the source in our analysis.

See "IV. Defining Individual Cases of Enemy Release and Inclusion Rules" for more detailed rules.

1. Paper Details

doi = Digital Object Identifier
filename = corresponding name of downloaded paper
title = title of paper

2. Inclusion Checklist

damagetype_accepted = compares a measure of enemy pressure between at least one 'enemy release' population and one 'control' population.
rawdata_available = raw data is available
mean_n_available = group average and number of replicates is available
enemy_release_focus = the paper looks at enemy release as a primary hypothesis
include_yn = source has passed the damagetype check and has either raw or mean+n data available

notes = any notes you may have
experiment_type = whether paper contains biogeographical or community studies (or both)

------------------------------------------
III. Calculating enemy release (data_main)
-------------------------------------------
This table is a list of each case of enemy release and supporting information.
Note that each source paper can have more than one or case associated with it 
under these rules.

1. Source Information
source_doi = DOI of source
source_filename = filename of source (also a study identifier)

2. Case Information
study_species = species name of the non-native population
system_type = general system (t = terrestrial, m = marine, f = freshwater)
species_type1 = taxa or type of study species
species_type2 = more specific taxa or type of study species
enemy_type = taxa of enemy or relationship that the enemy has with the study species 
(a = animal (excluding invertebrate herbivores*), p = plant, i = invertebrate herbivore, o = other (eg. fungi, nematodes, general microbes))

response = whether the measure is based on study species fitness, damage or diversity
damage = whether a positive values is related to greater(1) or less (-1) enemy effect.
effect_type = type of enemy damage as stated in source

n1 = number of samples of introduced pop
mean1 = mean of introduced pop
var1 = (pooled & weighted) sample variance of introduced pop
sd1 = standard deviation of introduced pop*
se1 = standard error of introduced pop*
ci1 = 95% confidence interval of introduced pop*
 
n2 = number of samples of control pop (if there are multiple control populations, it is the sum of them)
mean2 = mean of control pop
var2 = (pooled & weighted) sample variance of control pop
sd2 = standard deviation of control pop*
se2 = standard error of control pop*
ci2 = 95% confidence interval of control pop*

*if available, can be used as inputs to help calculate the variance

3.Other Useful Information

experiment_type = biogeographical or community comparison (b/c)
lab or field = whether experiment is in the field/manipulated in the lab (field/lab)
enemy = if the experiment tests the effect of a specific enemy, it is named here.
congener = if the experiment compares the study species with a specific congener, it is named here
notes = generic field to write notes in

Note that location only refers to the source of the introduced population.
location_lat = latitude (in decimal degrees N)
location_long = longitude (in decimal degrees E)
datum = datum used. If none given, it is inferred from publishing date+corresponding author address. 
Usually WGS84 unless paper is older or a local datum is used (we manually check through google earth if it looks questionable).

If no location provided but a placename is used, google maps is used with consideration for author affilations 
(and if required, gbif organism distrubution) to double check.
If multiple locations were tested from one population, geographic midpoint of all sampling locations was calculated.
time_start = start of introduced population records.
time_end = end of introduced population records. If only one time given, end is left blank.

If none provided, year of experiment end is inferred as 6 months before recieved publication date 
and then the start is estimated from the methods.

----------------------------------------------------------------------
IV. Defining Individual Cases of Enemy Release and Inclusion Rules
----------------------------------------------------------------------
1. Defining enemy release
Enemy release will be defined in this study by the difference in fitness (caused by enemy damage) between two groups, 
one that may have enemy release and a control that is not experiencing enemy release. 

It is generally tested by either:

#The Biogeographic Method#
Difference in enemy damage between 2+ populations of a single species; one study population in an introduced range and 1+ populations in a native range.

#The Community Method#
Difference in enemy damage between 2+ species; one study species population in an introduced range and 1+ cogeneric native species populations in the same range.

2. Defining one case
Each row is one measurement of enemy release.
Each row contains one case of an introduced population of a species within a specific space and time
(or a group of probably interbreeding populations),
compared with some theoretically non-enemy released control populations as a baseline,
with a quantitiative measurement of the enemy damage experienced by that population by a specific source.

2a. Enemy
Enemy damage will be grouped as one case if the enemies are primarily affected by similar defences.
Thus, if a paper records seperate values for different distinct groups or pairs of organism that caused detectably different types of enemy damage (eg. fungi vs insect),
each group will have its own record. However, enemy damage of the same type will be grouped as replicates of a single form of enemy damage, 
as the defence mechanisms against both organisms will significantly overlap (eg. two different specialised fungi).

2b. Measurements Taken
If multiple effects are available, the more direct measurement of detriment to fitness was used (ie. dmg>count>div, seed_dmg>flower_dmg>leaf_dmg).

2c. Distance 
If sites on a single paper are stated as the same population of eachother, measurements will be considered replicates of the same case of enemy release.

2d. Time
Repeat measurements in the same place throughout time are assumed to be one case unless it is stated that local extinction and re-introduction has occured between measurements.

2e. Replicates
Replicates (n) are defined by the original paper's experiment design. 
While number of plots is ideal to suit our study, the standard error measurements are linked to n from the source study design.

2f. Combination of experimental groups
When combining multiple groups is required to fit existing data into our introduced vs control structure:
Sample means will be weighted (by number of samples of each group) and pooled
Standard errors of the difference between means is calculated in accordance with https://doi.org/10.1007/978-94-011-7241-7_15 (as sqrt(SE1^2+SE2^2))

2g. Other Inclusion Rules
The following rules have been applied:
i. When a study applies multiple tests to its populations and some tests to only a subset of these populations, 
whatever test that provides the closest approximation of pure enemy effects will be chosen 
(as long as a introduced and comparison control population remain). 

ii. When both biogeographical and community tests have been performed on the same case,
biogeographical tests will be favoured.

iii. Each study was screened for retractions using the retractionwatch database (http://retractiondatabase.org/RetractionSearch.aspx?).
