## Supplementary File 3 Additional information for "How strong is enemy release? A systematic compilation across taxa and approaches"

**Supplementary File 3: Methods including search string, PRISMA documentation and references for enemy release dataset.**

Supplementary methods for “How strong is enemy release? A systematic compilation across taxa and approaches.” by K. Zeng, J. Schembri, E. Slavich, A. T. Moles.

Includes the following:

**Supplementary Methods A**: Initial Search

**Supplementary Methods B:** PRISMA Documentation

**Supplementary Methods C:** Reference list used to produce Enemy release literature synthesis dataset

**Supplementary Graph D:** Support for the enemy release hypothesis by record vs by study.

**Supplementary Methods A**: Initial Search

I performed a literature search on two of the largest scientific literature databases in ecology (WOS and SCOPUS). “Enemy Release” was searched on both SCOPUS and ISI WOS.

On ISI Web of Science, citation reports were saved in sets of 500 for a total of 1311 (accessed 25/05/2020). The exact search was as follows:

Query: (ALL = ("enemy" AND "release")) OR ALL=("enemy release")

Date: 1/1/1900 to 25/5/2020

Results: 1311

Format: BibTeX files (wos500.bib, wos1000.bib, wos1311.bib)

On Scopus, citation reports were saved in a single file of n (accessed 25/5/2020). The exact search was as follows:

Query: TITLE-ABS-KEY (“enemy" AND "release" )

Date: 25/5/2020

Results: 1857

Format: Research Information Systems file (scopus.ris)

Additional papers were identified after preliminary analysis by examining reference lists in original articles and prominent reviews.

**Supplementary Methods B:** PRISMA Documentation

Records identified from* databases:

Scopus (n = 1311)

Web of Science (n = 1857)

Records removed *before screening*:

Duplicate records removed (n = 2347)

Records marked as ineligible by automation tools (n = 0)

**Identification**

Records screened

(n = 821)

Records excluded

Unrelated/ Review/ Compilation (n = 193)

(n = )

**Screening**

Reports sought for retrieval

Automatic download (n = 275)

Manual download (n = 348)

Reports not retrieved

Record not available (n = 193)

Reports assessed for eligibility

(n = 628)

Reports excluded:

Record has no useable data (n = 405)

Record excluded upon closer inspection (23) (see below)

Record format unsuitable for our comparison (n = 2)

Locality unclear (n = 1)

Record is salami published so duplicate copies of data excluded (n = 20)

Studies included in review

(n = 225)

Reports of included studies

(n = 714)

**Included**

**Biogeographical Enemy Release (n = 380)**

**Community Enemy Release**

**(n = 311)**

Studies included in review

(n = 223)

Reports of included studies

(n = 691)

| **Section and Topic** | **Item #** | **Checklist item** | **Location where item is reported** |
| --- | --- | --- | --- |
| **TITLE** | | |  |
| Title | 1 | Identify the report as a systematic review. | Methods |
| **ABSTRACT** | | |  |
| Abstract | 2 | See the PRISMA 2020 for Abstracts checklist. |  |
| **INTRODUCTION** | | |  |
| Rationale | 3 | Describe the rationale for the review in the context of existing knowledge. | Introduction |
| Objectives | 4 | Provide an explicit statement of the objective(s) or question(s) the review addresses. | Introduction |
| **METHODS** | | |  |
| Eligibility criteria | 5 | Specify the inclusion and exclusion criteria for the review and how studies were grouped for the syntheses. | Methods |
| Information sources | 6 | Specify all databases, registers, websites, organisations, reference lists and other sources searched or consulted to identify studies. Specify the date when each source was last searched or consulted. | Methods |
| Search strategy | 7 | Present the full search strategies for all databases, registers and websites, including any filters and limits used. | Methods, SI |
| Selection process | 8 | Specify the methods used to decide whether a study met the inclusion criteria of the review, including how many reviewers screened each record and each report retrieved, whether they worked independently, and if applicable, details of automation tools used in the process. | Methods, SI |
| Data collection process | 9 | Specify the methods used to collect data from reports, including how many reviewers collected data from each report, whether they worked independently, any processes for obtaining or confirming data from study investigators, and if applicable, details of automation tools used in the process. | Methods, SI |
| Data items | 10a | List and define all outcomes for which data were sought. Specify whether all results that were compatible with each outcome domain in each study were sought (e.g. for all measures, time points, analyses), and if not, the methods used to decide which results to collect. | SI |
|  | 10b | List and define all other variables for which data were sought (e.g. participant and intervention characteristics, funding sources). Describe any assumptions made about any missing or unclear information. | SI |
| Study risk of bias assessment | 11 | Specify the methods used to assess risk of bias in the included studies, including details of the tool(s) used, how many reviewers assessed each study and whether they worked independently, and if applicable, details of automation tools used in the process. | Methods |
| Effect measures | 12 | Specify for each outcome the effect measure(s) (e.g. risk ratio, mean difference) used in the synthesis or presentation of results. | Methods |
| Synthesis methods | 13a | Describe the processes used to decide which studies were eligible for each synthesis (e.g. tabulating the study intervention characteristics and comparing against the planned groups for each synthesis (item #5)). | Methods |
|  | 13b | Describe any methods required to prepare the data for presentation or synthesis, such as handling of missing summary statistics, or data conversions. | Methods,  Github |
|  | 13c | Describe any methods used to tabulate or visually display results of individual studies and syntheses. | Github |
|  | 13d | Describe any methods used to synthesize results and provide a rationale for the choice(s). If meta-analysis was performed, describe the model(s), method(s) to identify the presence and extent of statistical heterogeneity, and software package(s) used. | Github |
|  | 13e | Describe any methods used to explore possible causes of heterogeneity among study results (e.g. subgroup analysis, meta-regression). | NA |
|  | 13f | Describe any sensitivity analyses conducted to assess robustness of the synthesized results. | NA |
| Reporting bias assessment | 14 | Describe any methods used to assess risk of bias due to missing results in a synthesis (arising from reporting biases). | NA |
| Certainty assessment | 15 | Describe any methods used to assess certainty (or confidence) in the body of evidence for an outcome. | NA |
| **RESULTS** | | |  |
| Study selection | 16a | Describe the results of the search and selection process, from the number of records identified in the search to the number of studies included in the review, ideally using a flow diagram. | SI |
|  | 16b | Cite studies that might appear to meet the inclusion criteria, but which were excluded, and explain why they were excluded. | Methods |
| Study characteristics | 17 | Cite each included study and present its characteristics. | SI |
| Risk of bias in studies | 18 | Present assessments of risk of bias for each included study. | NA |
| Results of individual studies | 19 | For all outcomes, present, for each study: (a) summary statistics for each group (where appropriate) and (b) an effect estimate and its precision (e.g. confidence/credible interval), ideally using structured tables or plots. | Methods, SI |
| Results of syntheses | 20a | For each synthesis, briefly summarise the characteristics and risk of bias among contributing studies. | Methods |
|  | 20b | Present results of all statistical syntheses conducted. If meta-analysis was done, present for each the summary estimate and its precision (e.g. confidence/credible interval) and measures of statistical heterogeneity. If comparing groups, describe the direction of the effect. | Results |
|  | 20c | Present results of all investigations of possible causes of heterogeneity among study results. | NA |
|  | 20d | Present results of all sensitivity analyses conducted to assess the robustness of the synthesized results. | NA |
| Reporting biases | 21 | Present assessments of risk of bias due to missing results (arising from reporting biases) for each synthesis assessed. | NA |
| Certainty of evidence | 22 | Present assessments of certainty (or confidence) in the body of evidence for each outcome assessed. | Results |
| **DISCUSSION** | | |  |
| Discussion | 23a | Provide a general interpretation of the results in the context of other evidence. | Discussion |
|  | 23b | Discuss any limitations of the evidence included in the review. | Discussion |
|  | 23c | Discuss any limitations of the review processes used. | Discussion |
|  | 23d | Discuss implications of the results for practice, policy, and future research. | Discussion |
| **OTHER INFORMATION** | | |  |
| Registration and protocol | 24a | Provide registration information for the review, including register name and registration number, or state that the review was not registered. | NA |
|  | 24b | Indicate where the review protocol can be accessed, or state that a protocol was not prepared. | NA |
|  | 24c | Describe and explain any amendments to information provided at registration or in the protocol. | NA |
| Support | 25 | Describe sources of financial or non-financial support for the review, and the role of the funders or sponsors in the review. | Acknowledgements |
| Competing interests | 26 | Declare any competing interests of review authors. | Acknowledgements |
| Availability of data, code and other materials | 27 | Report which of the following are publicly available and where they can be found: template data collection forms; data extracted from included studies; data used for all analyses; analytic code; any other materials used in the review. | Methods |

*From:*  Page MJ, McKenzie JE, Bossuyt PM, Boutron I, Hoffmann TC, Mulrow CD, et al. The PRISMA 2020 statement: an updated guideline for reporting systematic reviews. BMJ 2021;372:n71. doi: 10.1136/bmj.n71

For more information, visit: <http://www.prisma-statement.org/>

**Supplementary Methods C:** Reference list used to produce Enemy release literature synthesis dataset.

Abhilasha D, Joshi J (2009) Enhanced fitness due to higher fecundity, increased defence against a specialist and tolerance towards a generalist herbivore in an invasive annual plant. Journal of Plant Ecology 2: 77–86. https://doi.org/10.1093/jpe/rtp008

Adams JM, Fang W, Callaway RM, Cipollini D, and EN (2008) A cross-continental test of the Enemy Release Hypothesis: leaf herbivory on Acer platanoides (L.) is three times lower in North America than in its native Europe. Biological Invasions 11: 1005–1016. https://doi.org/10.1007/s10530-008-9312-4

Agrawal AA, Kotanen PM (2003) Herbivores and the success of exotic plants: a phylogenetically controlled experiment. Ecology Letters 6: 712–715. https://doi.org/10.1046/j.1461-0248.2003.00498.x

Alba C, Hufbauer R (2012) Exploring the potential for climatic factors, herbivory, and co-occurring vegetation to shape performance in native and introduced populations of Verbascum thapsus. Biological Invasions 14: 2505–2518. https://doi.org/10.1007/s10530-012-0247-4

Aldorfová A, Münzbergová Z (2019) Conditions of plant cultivation affect the differences in intraspecific plant-soil feedback between invasive and native dominants. Flora 261: 151492. https://doi.org/10.1016/j.flora.2019.151492

Allen WJ, Young RE, Bhattarai GP, Croy JR, Lambert AM, Meyerson LA, Cronin JT (2015) Multitrophic enemy escape of invasive Phragmites australis and its introduced herbivores in North America. Biological Invasions 17: 3419–3432. https://doi.org/10.1007/s10530-015-0968-2

Allen WJ, DeVries AE, Bologna NJ, Bickford WA, Kowalski KP, Meyerson LA, Cronin JT (2020) Intraspecific and biogeographical variation in foliar fungal communities and pathogen damage of native and invasive Phragmites australis. Moles A (Ed.). Global Ecology and Biogeography 29: 1199–1211. https://doi.org/10.1111/geb.13097

Ando Y, Utsumi S, Ohgushi T (2010) Community structure of insect herbivores on introduced and native Solidago plants in Japan. Entomologia Experimentalis et Applicata 136: 174–183. https://doi.org/10.1111/j.1570-7458.2010.01017.x

Andonian K, Hierro JL, Khetsuriani L, Becerra P, Janoyan G, Villarreal D, Cavieres L, Fox LR, Callaway RM (2011) Range-Expanding Populations of a Globally Introduced Weed Experience Negative Plant-Soil Feedbacks. Wright J (Ed.). PLoS ONE 6: e20117. https://doi.org/10.1371/journal.pone.0020117

Antonini Y, Lobato DNC, Norte AC, Ramos JA, Moreira P de A, Braga EM (2019) Patterns of avian malaria in tropical and temperate environments: testing the “The enemy release hypothesis.” Biota Neotropica 19. https://doi.org/10.1590/1676-0611-bn-2018-0716

Arundell K, Dunn A, Alexander J, Shearman R, Archer N, Ironside JE (2014) Enemy release and genetic founder effects in invasive killer shrimp populations of Great Britain. Biological Invasions 17: 1439–1451. https://doi.org/10.1007/s10530-014-0806-y

Avanesyan A, Culley TM (2015) Herbivory of native and exotic North-American prairie grasses by nymph Melanoplus grasshoppers. Plant Ecology 216: 451–464. https://doi.org/10.1007/s11258-015-0449-9

Bailly J, Garnier S, Khimoun A, Arnoux E, Eraud C, Goret J-Y, Luglia T, Gaucher P, Faivre B (2016) Reduced inflammation in expanding populations of a neotropical bird species. Ecology and Evolution 6: 7511–7521. https://doi.org/10.1002/ece3.2486

Beest M te, Stevens N, Olff H, Putten WH van der (2009) Plant-soil feedback induces shifts in biomass allocation in the invasive plant *Chromolaena odorata.* Journal of Ecology 97: 1281–1290. https://doi.org/10.1111/j.1365-2745.2009.01574.x

Bennett AE, Strauss SY (2012) Response to soil biota by native, introduced non-pest, and pest grass species: is responsiveness a mechanism for invasion? Biological Invasions 15: 1343–1353. https://doi.org/10.1007/s10530-012-0371-1

Blaisdell GK, Roy BA (2013) Two tests of enemy release of commonly co-occurring bunchgrasses native in Europe and introduced in the United States. Biological Invasions 16: 833–842. https://doi.org/10.1007/s10530-013-0541-9

Bodawatta KH, Clark C, Hedrick A, Hood A, Smith BH (2019) Comparative Analyses of Herbivory Rates and Leaf Phenology in Invasive and Native Shrubs in an East-Central Indiana Forest1. The Journal of the Torrey Botanical Society 146: 48. https://doi.org/10.3159/torrey-d-18-00005

Boussellaa W, Neifar L, Goedknegt MA, Thieltges DW (2018) Lessepsian migration and parasitism: richness, prevalence and intensity of parasites in the invasive fish Sphyraena chrysotaenia compared to its native congener Sphyraena sphyraena in Tunisian coastal waters. PeerJ 6: e5558. https://doi.org/10.7717/peerj.5558

Brandenburger CR, Kim M, Slavich E, Meredith FL, Salminen J-P, Sherwin WB, Moles AT (2020) Evolution of defense and herbivory in introduced plants—Testing enemy release using a known source population, herbivore trials, and time since introduction. Ecology and Evolution 10: 5451–5463. https://doi.org/10.1002/ece3.6288

Britton-Simmons KH, Pister B, Sánchez I, Okamoto D (2010) Response of a native, herbivorous snail to the introduced seaweed Sargassum muticum. Hydrobiologia 661: 187–196. https://doi.org/10.1007/s10750-010-0523-1

BUSCHMANN H, EDWARDS PJ, DIETZ H (2005) Variation in growth pattern and response to slug damage among native and invasive provenances of four perennial Brassicaceae species. Journal of Ecology 93: 322–334. https://doi.org/10.1111/j.1365-2745.2005.00991.x

Cameron GN, Spencer SR (2010) Entomofauna of the Introduced Chinese Tallow Tree. The Southwestern Naturalist 55: 179–192. https://doi.org/10.1894/jc-28.1

Caño L, Escarré J, Vrieling K, Sans FX (2008) Palatability to a generalist herbivore, defence and growth of invasive and native Senecio species: testing the evolution of increased competitive ability hypothesis. Oecologia 159: 95–106. https://doi.org/10.1007/s00442-008-1182-z

Cardoso AC, Arenas F, Sousa-Pinto I, Barreiro A, Franco JN (2020) Sea urchin grazing preferences on native and non-native macroalgae. Ecological Indicators 111: 106046. https://doi.org/10.1016/j.ecolind.2019.106046

CARPENTER D, CAPPUCCINO N (2005) Herbivory, time since introduction and the invasiveness of exotic plants. Journal of Ecology 93: 315–321. https://doi.org/10.1111/j.1365-2745.2005.00973.x

Castells E, Morante M, Goula M, Pérez N, Dantart J, Escolà A (2013a) Herbivores on native and exotic Senecio plants: is host switching related to plant novelty and insect diet breadth under field conditions? Schonrogge K, Gange A (Eds). Insect Conservation and Diversity 7: 420–431. https://doi.org/10.1111/icad.12064

Castells E, Morante M, Blanco-Moreno JM, Sans FX, Vilatersana R, Blasco-Moreno A (2013b) Reduced seed predation after invasion supports enemy release in a broad biogeographical survey. Oecologia 173: 1397–1409. https://doi.org/10.1007/s00442-013-2718-4

Castillo G, Calahorra-Oliart A, Núñez-Farfán J, Valverde PL, Arroyo J, Cruz LL, Tapia-López R (2019) Selection on tropane alkaloids in native and non-native populations of Datura stramonium. Ecology and Evolution 9: 10176–10184. https://doi.org/10.1002/ece3.5520

Cincotta CL, Adams JM, Holzapfel C (2008) Testing the enemy release hypothesis: a comparison of foliar insect herbivory of the exotic Norway maple (Acer platanoides L.) and the native sugar maple (A. saccharum L.). Biological Invasions 11: 379–388. https://doi.org/10.1007/s10530-008-9255-9

Clark NJ, Olsson-Pons S, Ishtiaq F, Clegg SM (2015) Specialist enemies, generalist weapons and the potential spread of exotic pathogens: malaria parasites in a highly invasive bird. International Journal for Parasitology 45: 891–899. https://doi.org/10.1016/j.ijpara.2015.08.008

Cogni R (2009) Resistance to Plant Invasion? A Native Specialist Herbivore Shows Preference for and Higher Fitness on an Introduced Host. Biotropica 42: 188–193. https://doi.org/10.1111/j.1744-7429.2009.00570.x

Comont RF, Purse BV, Phillips W, Kunin WE, Hanson M, Lewis OT, Harrington R, Shortall CR, Rondoni G, Roy HE (2013) Escape from parasitism by the invasive alien ladybird, Harmonia axyridis. Leather SR, Sait S (Eds). Insect Conservation and Diversity 7: 334–342. https://doi.org/10.1111/icad.12060

CONNOR EF, FAETH SH, SIMBERLOFF D, OPLER PA (1980) Taxonomic isolation and the accumulation of herbivorous insects: a comparison of introduced and native trees. Ecological Entomology 5: 205–211.

Correia M, Montesinos D, French K, Rodríguez-Echeverría S (2016) Evidence for enemy release and increased seed production and size for two invasive Australian acacias. Mack R (Ed.). Journal of Ecology 104: 1391–1399. https://doi.org/10.1111/1365-2745.12612

Cripps MG, Schwarzländer M, McKenney JL, Hinz HL, Price WJ (2006) Biogeographical comparison of the arthropod herbivore communities associated with Lepidium draba in its native, expanded and introduced ranges. Journal of Biogeography 33: 2107–2119. https://doi.org/10.1111/j.1365-2699.2006.01560.x

Cripps MG, Hinz HL, McKenney JL, Price WJ, Schwarzländer M (2009) No evidence for an `evolution of increased competitive ability’ for the invasive Lepidium draba. Basic and Applied Ecology 10: 103–112. https://doi.org/10.1016/j.baae.2008.03.001

Cripps MG, Bourdôt GW, Saville DJ, Hinz HL, Fowler SV, Edwards GR (2011) Influence of insects and fungal pathogens on individual and population parameters of Cirsium arvense in its native and introduced ranges. Biological Invasions 13: 2739–2754. https://doi.org/10.1007/s10530-011-9944-7

Dang C, Montaudouin X de, Bald J, Jude F, Raymond N, Lanceleur L, Paul-Pont I, Caill-Milly N (2009) Testing the enemy release hypothesis: trematode parasites in the non-indigenous Manila clam Ruditapes philippinarum. Hydrobiologia 630: 139–148. https://doi.org/10.1007/s10750-009-9786-9

Dawson W, Bottini A, Fischer M, Kleunen M van, Knop E (2014) Little evidence for release from herbivores as a driver of plant invasiveness from a multi-species herbivore-removal experiment. Oikos 123: 1509–1518. https://doi.org/10.1111/oik.01485

Diagne C, Ribas A, Charbonnel N, Dalecky A, Tatard C, Gauthier P, Haukisalmi V, Fossati-Gaschignard O, Bâ K, Kane M, Niang Y, Diallo M, Sow A, Piry S, Sembène M, Brouat C (2016) Parasites and invasions: changes in gastrointestinal helminth assemblages in invasive and native rodents in Senegal. International Journal for Parasitology 46: 857–869. https://doi.org/10.1016/j.ijpara.2016.07.007

Dostál P, Allan E, Dawson W, Kleunen M van, Bartish I, Fischer M (2012) Enemy damage of exotic plant species is similar to that of natives and increases with productivity. Whitney K (Ed.). Journal of Ecology 101: 388–399. https://doi.org/10.1111/1365-2745.12037

Ebeling SK, Hensen I, Auge H (2007) The invasive shrub Buddleja davidii performs better in its introduced range. Diversity and Distributions 14: 225–233. https://doi.org/10.1111/j.1472-4642.2007.00422.x

Egan PA, Stevenson PC, Tiedeken EJ, Wright GA, Boylan F, Stout JC (2016) Plant toxin levels in nectar vary spatially across native and introduced populations. Phillips R (Ed.). Journal of Ecology 104: 1106–1115. https://doi.org/10.1111/1365-2745.12573

Elzinga JA, Bernasconi G (2009) Enhanced frugivory on invasive Silene latifolia in its native range due to increased oviposition. Journal of Ecology 97: 1010–1019. https://doi.org/10.1111/j.1365-2745.2009.01534.x

Endriss SB, Alba C, Norton AP, Pyšek P, Hufbauer RA (2017) Breakdown of a geographic cline explains high performance of introduced populations of a weedy invader. Rees M (Ed.). Journal of Ecology 106: 699–713. https://doi.org/10.1111/1365-2745.12845

Engelkes T, Wouters B, Bezemer TM, Harvey JA, Putten WH van der (2012) Contrasting patterns of herbivore and predator pressure on invasive and native plants. Basic and Applied Ecology 13: 725–734. https://doi.org/10.1016/j.baae.2012.10.005

Engelkes T, Morriën E, Verhoeven KJF, Bezemer TM, Biere A, Harvey JA, McIntyre LM, Tamis WLM, Putten WH van der (2008) Successful range-expanding plants experience less above-ground and below-ground enemy impact. Nature 456: 946–948. https://doi.org/10.1038/nature07474

Fairfield EA, Hutchings K, Gilroy DL, Kingma SA, Burke T, Komdeur J, Richardson DS (2016) The impact of conservation-driven translocations on blood parasite prevalence in the Seychelles warbler. Scientific Reports 6. https://doi.org/10.1038/srep29596

Fan S, Yu D, Liu C (2013) The Invasive Plant Alternanthera philoxeroides Was Suppressed More Intensively than Its Native Congener by a Native Generalist: Implications for the Biotic Resistance Hypothesis. Lamb EG (Ed.). PLoS ONE 8: e83619. https://doi.org/10.1371/journal.pone.0083619

Fan S, Yu H, Dong X, Wang L, Chen X, Yu D, Liu C (2016) Invasive plant Alternanthera philoxeroides suffers more severe herbivory pressure than native competitors in recipient communities. Scientific Reports 6. https://doi.org/10.1038/srep36542

Ferreras AE, Galetto L (2010) From seed production to seedling establishment: Important steps in an invasive process. Acta Oecologica 36: 211–218. https://doi.org/10.1016/j.actao.2009.12.005

Firmat C, Alibert P, Mutin G, Losseau M, Pariselle A, Sasal P (2016) A case of complete loss of gill parasites in the invasive cichlid Oreochromis mossambicus. Parasitology Research 115: 3657–3661. https://doi.org/10.1007/s00436-016-5168-1

Forslund H, Wikström SA, Pavia H (2010) Higher resistance to herbivory in introduced compared to native populations of a seaweed. Oecologia 164: 833–840. https://doi.org/10.1007/s00442-010-1767-1

Franks SJ, Pratt PD, Dray FA, Simms EL (2007) No evolution of increased competitive ability or decreased allocation to defense in Melaleuca quinquenervia since release from natural enemies. Biological Invasions 10: 455–466. https://doi.org/10.1007/s10530-007-9143-8

Gard B, Bretagnolle F, Dessaint F, Laitung B (2013) Invasive and native populations of common ragweed exhibit strong tolerance to foliar damage. Basic and Applied Ecology 14: 28–35. https://doi.org/10.1016/j.baae.2012.10.007

Gendron A, Marcogliese D (2016) Reduced survival of a native parasite in the invasive round goby: evidence for the dilution hypothesis? Aquatic Invasions 11: 189–198. https://doi.org/10.3391/ai.2016.11.2.08

Gendron AD, Marcogliese DJ, Thomas M (2011) Invasive species are less parasitized than native competitors, but for how long? The case of the round goby in the Great Lakes-St. Lawrence Basin. Biological Invasions 14: 367–384. https://doi.org/10.1007/s10530-011-0083-y

Genton BJ, Kotanen PM, Cheptou P-O, Adolphe C, Shykoff JA (2005) Enemy release but no evolutionary loss of defence in a plant invasion: an inter-continental reciprocal transplant experiment. Oecologia 146: 404–414. https://doi.org/10.1007/s00442-005-0234-x

Goedknegt A, Havermans J, Waser A, Luttikhuizen P, Velilla E, Camphuysen K, Meer J van der, Thieltges D (2017) Cross-species comparison of parasite richness, prevalence, and intensity in a native compared to two invasive brachyuran crabs. Aquatic Invasions 12: 201–212. https://doi.org/10.3391/ai.2017.12.2.08

Goergen E, Daehler CC (2001) Inflorescence Damage by Insects and Fungi in Native Pili Grass ( Heteropogon contortus ) versus Alien Foundation Grass ( Pennisetum setaceum ) in Hawai’i. Pacific Science 55: 129–136. https://doi.org/10.1353/psc.2001.0014

Gollan JR, Wright JT (2006) Limited grazing pressure by native herbivores on the invasive seaweed Caulerpa taxifolia in a temperate Australian estuary. Marine and Freshwater Research 57: 685. https://doi.org/10.1071/mf05253

González-Teuber M, Quiroz CL, Concha-Bloomfield I, Cavieres LA (2016) Enhanced fitness and greater herbivore resistance: implications for dandelion invasion in an alpine habitat. Biological Invasions 19: 647–653. https://doi.org/10.1007/s10530-016-1309-9

Grunsven RHAV, Bos F, Ripley BS, Suehs CM, Veenendaal EM (2009) Release from soil pathogens plays an important role in the success of invasive Carpobrotus in the Mediterranean. South African Journal of Botany 75: 172–175. https://doi.org/10.1016/j.sajb.2008.09.003

GRUNSVEN RHAV, PUTTEN WHVD, BEZEMER TM, TAMIS WLM, BERENDSE F, VEENENDAAL EM (2007) Reduced plant-soil feedback of plant species expanding their range as compared to natives. Journal of Ecology 95: 1050–1057. https://doi.org/10.1111/j.1365-2745.2007.01282.x

Gruntman M, Segev U, Glauser G, Tielbörger K (2016) Evolution of plant defences along an invasion chronosequence: defence is lost due to enemy release – but not forever. Bartomeus I (Ed.). Journal of Ecology 105: 255–264. https://doi.org/10.1111/1365-2745.12660

Gundale MJ, Almeida JP, Wallander H, Wardle DA, Kardol P, Nilsson M-C, Fajardo A, Pauchard A, Peltzer DA, Ruotsalainen S, Mason B, Rosenstock N (2016) Differences in endophyte communities of introduced trees depend on the phylogenetic relatedness of the receiving forest. Austin A (Ed.). Journal of Ecology 104: 1219–1232. https://doi.org/10.1111/1365-2745.12595

Halbritter AH, Carroll GC, Güsewell S, Roy BA (2012) Testing assumptions of the enemy release hypothesis: generalist versus specialist enemies of the grass Brachypodium sylvaticum. Mycologia 104: 34–44. https://doi.org/10.3852/11-071

Hammann M, Wang G, Rickert E, Boo SM, Weinberger F (2013) Invasion success of the seaweed Gracilaria vermiculophylla correlates with low palatibility. Marine Ecology Progress Series 486: 93–103. https://doi.org/10.3354/meps10361

Han X, Dendy SP, Garrett KA, Fang L, Smith MD (2008) Comparison of damage to native and exotic tallgrass prairie plants by natural enemies. Plant Ecology 198: 197–210. https://doi.org/10.1007/s11258-008-9395-0

HANLEY ME (2012) Seedling defoliation, plant growth and flowering potential in native- and invasive-range Plantago lanceolata populations. Weed Research 52: 252–259. https://doi.org/10.1111/j.1365-3180.2012.00910.x

HANSEN SO, HATTENDORF J, WITTENBERG R, REZNIK SY, NIELSEN C, RAVN HP, NENTWIG W (2006) Phytophagous insects of giant hogweed Heracleum mantegazzianum (Apiaceae) in invaded areas of Europe and in its native area of the Caucasus. European Journal of Entomology 103: 387–395. https://doi.org/10.14411/eje.2006.052

Hartley MK, Rogers WE, Siemann E (2010) Comparisons of arthropod assemblages on an invasive and native trees: abundance, diversity and damage. Arthropod-Plant Interactions 4: 237–245. https://doi.org/10.1007/s11829-010-9105-4

Harvey KJ, Nipperess DA, Britton DR, Hughes L (2015) Comparison of invertebrate herbivores on native and non-native Senecio species: Implications for the enemy release hypothesis. Austral Ecology 40: 503–514. https://doi.org/10.1111/aec.12216

Hazell SP, Vel T, Fellowes MDE (2007) The role of exotic plants in the invasion of Seychelles by the polyphagous insect Aleurodicus dispersus: a phylogenetically controlled analysis. Biological Invasions 10: 169–175. https://doi.org/10.1007/s10530-007-9120-2

Hill SB, Kotanen PM (2010) Phylogenetically structured damage to Asteraceae: susceptibility of native and exotic species to foliar herbivores. Biological Invasions 12: 3333–3342. https://doi.org/10.1007/s10530-010-9726-7

Hintz WD, Schuler MS, Jones DK, Coldsnow KD, Stoler AB, Relyea RA (2019) Nutrients influence the multi-trophic impacts of an invasive species unaffected by native competitors or predators. Science of The Total Environment 694: 133704. https://doi.org/10.1016/j.scitotenv.2019.133704

Hinz HL, Schwarzländer M, McKenney JL, Cripps MG, Harmon B, Price WJ (2012) Biogeographical comparison of the invasive Lepidium draba in its native, expanded and introduced ranges. Biological Invasions 14: 1999–2016. https://doi.org/10.1007/s10530-012-0207-z

Hopper JV, Kuris AM, Lorda J, Simmonds SE, White C, Hechinger RF (2014) Reduced parasite diversity and abundance in a marine whelk in its expanded geographical range. Veech J (Ed.). Journal of Biogeography 41: 1674–1684. https://doi.org/10.1111/jbi.12329

Hornoy B, Tarayre M, Hervé M, Gigord L, Atlan A (2011) Invasive Plants and Enemy Release: Evolution of Trait Means and Trait Correlations in Ulex europaeus. Moora M (Ed.). PLoS ONE 6: e26275. https://doi.org/10.1371/journal.pone.0026275

Huang W, Ding J (2015) Effects of generalist herbivory on resistance and resource allocation by the invasive plant, Phytolacca americana. Insect Science 23: 191–199. https://doi.org/10.1111/1744-7917.12244

Ishtiaq F, Beadell JS, Baker AJ, Rahmani AR, Jhala YV, Fleischer RC (2005) Prevalence and evolutionary relationships of haematozoan parasites in native versus introduced populations of common myna Acridotheres tristis. Proceedings of the Royal Society B: Biological Sciences 273: 587–594. https://doi.org/10.1098/rspb.2005.3313

Jack CN, Friesen ML (2019) Rapid evolution of Medicago polymorpha during invasion shifts interactions with the soybean looper. Ecology and Evolution 9: 10522–10533. https://doi.org/10.1002/ece3.5572

Jones CM, Brown MJF (2014) Parasites and genetic diversity in an invasive bumblebee. Ings T (Ed.). Journal of Animal Ecology 83: 1428–1440. https://doi.org/10.1111/1365-2656.12235

Joshi S, Tielbörger K (2012) Response to enemies in the invasive plant Lythrum salicaria is genetically determined. Annals of Botany 110: 1403–1410. https://doi.org/10.1093/aob/mcs076

Katz DSW, Ibáñez I (2016) Differences in biotic interactions across range edges have only minor effects on plant performance. Heard M (Ed.). Journal of Ecology 105: 321–331. https://doi.org/10.1111/1365-2745.12675

Kimball S, Gremer JR, Barron-Gafford GA, Angert AL, Huxman TE, Venable DL (2014) High water-use efficiency and growth contribute to success of non-native Erodium cicutarium in a Sonoran Desert winter annual community. Conservation Physiology 2: cou006–cou006. https://doi.org/10.1093/conphys/cou006

Kirichenko N, Kenis M (2016) Using a botanical garden to assess factors influencing the colonization of exotic woody plants by phyllophagous insects. Oecologia 182: 243–252. https://doi.org/10.1007/s00442-016-3645-y

Knapp LB, Fownes JH, Harrington RA (2008) Variable effects of large mammal herbivory on three non-native versus three native woody plants. Forest Ecology and Management 255: 92–98. https://doi.org/10.1016/j.foreco.2007.08.023

Knevel IC, Lans T, Menting FBJ, Hertling UM, Putten WH van der (2004) Release from native root herbivores and biotic resistance by soil pathogens in a new habitat both affect the alien Ammophila arenaria in South Africa. Oecologia 141: 502–510. https://doi.org/10.1007/s00442-004-1662-8

Korell L, Schädler M, Brandl R, Schreiter S, Auge H (2019) Release from Above- and Belowground Insect Herbivory Mediates Invasion Dynamics and Impact of an Exotic Plant. Plants 8: 544. https://doi.org/10.3390/plants8120544

Korell L, Stein C, Hensen I, Bruelheide H, Suding KN, Auge H (2016) Stronger effect of gastropods than rodents on seedling establishment, irrespective of exotic or native plant species origin. Oikos 125: 1467–1477. https://doi.org/10.1111/oik.02696

Krakau M, Thieltges DW, Reise K (2006) Native Parasites Adopt Introduced Bivalves of the North Sea. Biological Invasions 8: 919–925. https://doi.org/10.1007/s10530-005-4734-8

Kruse J, Pautasso M, Aas G (2016) A test of the enemy release hypothesis for plants in the Ecological-Botanical Gardens, Bayreuth, using data on plant parasitic microfungi. Nova Hedwigia 103: 239–249. https://doi.org/10.1127/nova_hedwigia/2016/0348

Kumschick S, Hufbauer RA, Alba C, Blumenthal DM (2013) Evolution of fast-growing and more resistant phenotypes in introduced common mullein (Verbascum thapsus). Prentice HC (Ed.). Journal of Ecology 101: 378–387. https://doi.org/10.1111/1365-2745.12044

Kwong RM, Sagliocco JL, Harms NE, Butler KL, Martin GD, Green PT (2019) Could enemy release explain invasion success of Sagittaria platyphylla in Australia and South Africa? Aquatic Botany 153: 67–72. https://doi.org/10.1016/j.aquabot.2018.11.011

Lacerda ACF, Takemoto RM, Poulin R, Pavanelli GC (2012) Parasites of the fish Cichla piquiti (Cichlidae) in native and invaded Brazilian basins: release not from the enemy, but from its effects. Parasitology Research 112: 279–288. https://doi.org/10.1007/s00436-012-3135-z

Lakeman-Fraser P, Ewers RM (2012) Enemy release promotes range expansion in a host plant. Oecologia 172: 1203–1212. https://doi.org/10.1007/s00442-012-2555-x

Lapointe M, Brisson J (2012) A Comparison of Invasive Acer platanoides and Native A. saccharum First-Year Seedlings: Growth, Biomass Distribution and the Influence of Ecological Factors in a Forest Understory. Forests 3: 190–206. https://doi.org/10.3390/f3020190

Larson MD (2018) Range Expansion and Parasitism in the Nonnative Snail Radix auricularia. Western North American Naturalist 78: 112. https://doi.org/10.3398/064.078.0112

Leishman MR, Cooke J, Richardson DM (2014) Evidence for shifts to faster growth strategies in the new ranges of invasive alien plants. Newman J (Ed.). Journal of Ecology 102: 1451–1461. https://doi.org/10.1111/1365-2745.12318

Lester PJ, Gruber MAM, Brenton-Rule EC, Archer M, Corley JC, Dvořák L, Masciocchi M, Oystaeyen AV (2014) Determining the origin of invasions and demonstrating a lack of enemy release from microsporidian pathogens in common wasps (Vespula vulgaris). Roura-Pascual N (Ed.). Diversity and Distributions 20: 964–974. https://doi.org/10.1111/ddi.12223

Lester PJ, Bosch PJ, Gruber MAM, Kapp EA, Peng L, Brenton-Rule EC, Buchanan J, Stanislawek WL, Archer M, Corley JC, Masciocchi M, Oystaeyen AV, Wenseleers T (2015) No Evidence of Enemy Release in Pathogen and Microbial Communities of Common Wasps (Vespula vulgaris) in Their Native and Introduced Range. Lee B-L (Ed.). PLOS ONE 10: e0121358. https://doi.org/10.1371/journal.pone.0121358

Lewis KC, Bazzaz FA, Liao Q, Orians CM (2006) Geographic patterns of herbivory and resource allocation to defense, growth, and reproduction in an invasive biennial, Alliaria petiolata. Oecologia 148: 384–395. https://doi.org/10.1007/s00442-006-0380-9

Li Y, Ke Z, Wang S, Smith GR, Liu X (2011) An Exotic Species Is the Favorite Prey of a Native Enemy. Hayward M (Ed.). PLoS ONE 6: e24299. https://doi.org/10.1371/journal.pone.0024299

Liao Z-Y, Zheng Y-L, Lei Y-B, Feng Y-L (2013) Evolutionary increases in defense during a biological invasion. Oecologia 174: 1205–1214. https://doi.org/10.1007/s00442-013-2852-z

Lieurance D, Cipollini D (2013) Exotic Lonicera species both escape and resist specialist and generalist herbivores in the introduced range in North America. Biological Invasions 15: 1713–1724. https://doi.org/10.1007/s10530-012-0402-y

Lieurance D, Chakraborty S, Whitehead SR, Powell JR, Bonello P, Bowers MD, Cipollini D (2015) Comparative Herbivory Rates and Secondary Metabolite Profiles in the Leaves of Native and Non-Native Lonicera Species. Journal of Chemical Ecology 41: 1069–1079. https://doi.org/10.1007/s10886-015-0648-9

Lima M, Simpson L, Fecchio A, Kyaw C (2010) Low prevalence of haemosporidian parasites in the introduced house sparrow (Passer domesticus) in Brazil. Acta Parasitologica 55. https://doi.org/10.2478/s11686-010-0055-x

Liu H, Stiling P, Pemberton RW (2007) Does enemy release matter for invasive plants? evidence from a comparison of insect herbivore damage among invasive, non-invasive and native congeners. Biological Invasions 9: 773–781. https://doi.org/10.1007/s10530-006-9074-9

Lombardero MJ, Vázquez-Mejuto P, Ayres MP (2008) ROLE OF PLANT ENEMIES IN THE FORESTRY OF INDIGENOUS VS. NONINDIGENOUS PINES. Ecological Applications 18: 1171–1181. https://doi.org/10.1890/07-1048.1

Lopez VM, Hoddle MS (2013) Mortality factors affecting Agrilus auroguttatus Schaeffer (Coleoptera: Buprestidae) eggs in the native and invaded ranges. Biological Control 67: 143–148. https://doi.org/10.1016/j.biocontrol.2013.07.006

Lu X, He M, Tang S, Wu Y, Shao X, Wei H, Siemann E, Ding J (2019) Herbivory may promote a non-native plant invasion at low but not high latitudes. Annals of Botany 124: 819–827. https://doi.org/10.1093/aob/mcz121

Lucero JE, Callaway RM (2018) Native granivores reduce the establishment of native grasses but not invasive Bromus tectorum. Biological Invasions 20: 3491–3497. https://doi.org/10.1007/s10530-018-1789-x

Lucero JE, Schaffner U, Asadi G, Bagheri A, Rajabov T, Callaway RM (2019) Enemy release from the effects of generalist granivores can facilitate Bromus tectorum invasion in the Great Basin Desert. Ecology and Evolution 9: 8490–8499. https://doi.org/10.1002/ece3.5314

Lu-Irving P, Harenčár JG, Sounart H, Welles SR, Swope SM, Baltrus DA, Dlugosch KM (2019) Native and Invading Yellow Starthistle (Centaurea solstitialis) Microbiomes Differ in Composition and Diversity of Bacteria. McMahon K (Ed.). mSphere 4. https://doi.org/10.1128/msphere.00088-19

Macel M, Dostálek T, Esch S, Bucharová A, Dam NM van, Tielbörger K, Verhoeven KJF, Münzbergová Z (2017) Evolutionary responses to climate change in a range expanding plant. Oecologia 184: 543–554. https://doi.org/10.1007/s00442-017-3864-x

MACLEOD CJ, DUNCAN RP, PARISH DMB, WRATTEN SD, HUBBARD SF (2005) Can increased niche opportunities and release from enemies explain the success of introduced Yellowhammer populations in New Zealand? Ibis 147: 598–607. https://doi.org/10.1111/j.1474-919x.2005.00445.x

Magro A, Ramon-Portugal F, Facon B, Ducamp C, Hemptinne J-L (2018) The evolution of chemical defenses along invasion routes: Harmonia axyridis Pallas (Coccinellidae: Coleoptera) as a case study. Ecology and Evolution 8: 8344–8353. https://doi.org/10.1002/ece3.4299

Maron JL, Vilà M, Arnason J (2004) LOSS OF ENEMY RESISTANCE AMONG INTRODUCED POPULATIONS OF ST. JOHN’S WORT (HYPERICUM PERFORATUM). Ecology 85: 3243–3253. https://doi.org/10.1890/04-0297

Maron JL, Klironomos J, Waller L, Callaway RM (2013) Invasive plants escape from suppressive soil biota at regional scales. Austin A (Ed.). Journal of Ecology 102: 19–27. https://doi.org/10.1111/1365-2745.12172

Marr SR, Mautz WJ, Hara AH (2007) Parasite loss and introduced species: a comparison of the parasites of the Puerto Rican tree frog, (Eleutherodactylus coqui), in its native and introduced ranges. Biological Invasions 10: 1289–1298. https://doi.org/10.1007/s10530-007-9203-0

Martin LB, Alam JL, Imboma T, Liebl AL (2010) Variation in inflammation as a correlate of range expansion in Kenyan house sparrows. Oecologia 164: 339–347. https://doi.org/10.1007/s00442-010-1654-9

Marzal A, Møller AP, Espinoza K, Morales S, Luján-Vega C, Cárdenas-Callirgos JM, Mendo L, Álvarez-Barrientos A, González-Blázquez M, García-Longoria L, Lope F de, Mendoza C, Iannacone J, Magallanes S (2018) Variation in malaria infection and immune defence in invasive and endemic house sparrows. Animal Conservation 21: 505–514. https://doi.org/10.1111/acv.12423

Marzal A, Ricklefs RE, Valkiūnas G, Albayrak T, Arriero E, Bonneaud C, Czirják GA, Ewen J, Hellgren O, Hořáková D, Iezhova TA, Jensen H, Križanauskien\.e A, Lima MR, Lope F de, Magnussen E, Martin LB, Møller AP, Palinauskas V, Pap PL, Pérez-Tris J, Sehgal RNM, Soler M, Szöll\Hosi E, Westerdahl H, Zetindjiev P, Bensch S (2011) Diversity, Loss, and Gain of Malaria Parasites in a Globally Invasive Bird. Fleischer RC (Ed.). PLoS ONE 6: e21905. https://doi.org/10.1371/journal.pone.0021905

Matter SF, Brzyski JR, Harrison CJ, Hyams S, Loo C, Loomis J, Lubbers HR, Seastrum L, Stamper TI, Stein AM, Stokes R, Wilkerson BS (2012) Invading from the garden? A comparison of leaf herbivory for exotic and native plants in natural and ornamental settings. Insect Science 19: 677–682. https://doi.org/10.1111/j.1744-7917.2012.01524.x

McGinn KJ, Putten WH van der, Hulme PE, Shelby N, Weser C, Duncan RP (2017) The influence of residence time and geographic extent on the strength of plant-soil feedbacks for naturalised Trifolium. Turnbull M (Ed.). Journal of Ecology 106: 207–217. https://doi.org/10.1111/1365-2745.12864

Meijer K, Zemel H, Chiba S, Smit C, Beukeboom LW, Schilthuizen M (2015) Phytophagous Insects on Native and Non-Native Host Plants: Combining the Community Approach and the Biogeographical Approach. Gaquerel E (Ed.). PLOS ONE 10: e0125607. https://doi.org/10.1371/journal.pone.0125607

Memmott J, Fowler SV, Paynter Q, Sheppard AW, Syrett P (2000) The invertebrate fauna on broom, Cytisus scoparius,in two native and two exotic habitats. Acta Oecologica 21: 213–222. https://doi.org/10.1016/s1146-609x(00)00124-7

MENÉNDEZ R, GONZÁLEZ-MEGÍAS A, LEWIS OT, SHAW MR, THOMAS CD (2008) Escape from natural enemies during climate-driven range expansion: a case study. Ecological Entomology 33: 413–421. https://doi.org/10.1111/j.1365-2311.2008.00985.x

Meyer G, Clare R, Weber E (2005) An experimental test of the evolution of increased competitive ability hypothesis in goldenrod, Solidago gigantea. Oecologia 144: 299–307. https://doi.org/10.1007/s00442-005-0046-z

Miller A, Inglis GJ, Poulin R (2008) Use of the introduced bivalve, Musculista senhousia, by generalist parasites of native New Zealand bivalves. New Zealand Journal of Marine and Freshwater Research 42: 143–151. https://doi.org/10.1080/00288330809509944

Mlynarek JJ (2015a) Testing the enemy release hypothesis in a native insect species with an expanding range. PeerJ 3: e1415. https://doi.org/10.7717/peerj.1415

Mlynarek JJ (2015b) Testing the enemy release hypothesis in a native insect species with an expanding range. PeerJ 3: e1415. https://doi.org/10.7717/peerj.1415

Monteiro CA, Engelen AH, Santos ROP (2009) Macro- and mesoherbivores prefer native seaweeds over the invasive brown seaweed Sargassum muticum: a potential regulating role on invasions. Marine Biology 156: 2505–2515. https://doi.org/10.1007/s00227-009-1275-1

Montti L, Ayup MM, Aragón R, Qi W, Ruan H, Fernández R, Casertano SA, Zou X (2016) Herbivory and the success of Ligustrum lucidum: evidence from a comparison between native and novel ranges. Australian Journal of Botany 64: 181. https://doi.org/10.1071/bt15232

Morriën E, Engelkes T, Putten WH van der (2011a) Additive effects of aboveground polyphagous herbivores and soil feedback in native and range-expanding exotic plants. Ecology 92: 1344–1352. https://doi.org/10.1890/10-1937.1

Morriën E, Duyts H, Putten WHV der (2011b) Effects of native and exotic range-expanding plant species on taxonomic and functional composition of nematodes in the soil food web. Oikos 121: 181–190. https://doi.org/10.1111/j.1600-0706.2011.19773.x

MORRISON JA, MAUCK K (2007) Experimental field comparison of native and non-native maple seedlings: natural enemies, ecophysiology, growth and survival. Journal of Ecology 95: 1036–1049. https://doi.org/10.1111/j.1365-2745.2007.01270.x

Müller G, Horstmeyer L, Rönneburg T, Kleunen M van, Dawson W (2016) Alien and native plant establishment in grassland communities is more strongly affected by disturbance than above- and below-ground enemies. Austin A (Ed.). Journal of Ecology 104: 1233–1242. https://doi.org/10.1111/1365-2745.12601

Naddafi R, Rudstam LG (2013) Predation on invasive zebra mussel, Dreissena polymorpha, by pumpkinseed sunfish, rusty crayfish, and round goby. Hydrobiologia 721: 107–115. https://doi.org/10.1007/s10750-013-1653-z

Najberek K, Solarz W, Chmura D (2017) Do local enemies attack alien and native Impatiens alike? Acta Societatis Botanicorum Poloniae 86. https://doi.org/10.5586/asbp.3562

Najberek K, Okarma H, Chmura D, Król W, Walusiak E, Solarz W (2019) Enemy pressure exerted on alien and native plants may differ between montane and lowland regions. Arthropod-Plant Interactions 14: 275–287. https://doi.org/10.1007/s11829-019-09736-6

Norghauer JM, Martin AR, Mycroft EE, James A, Thomas SC (2011) Island Invasion by a Threatened Tree Species: Evidence for Natural Enemy Release of Mahogany (Swietenia macrophylla) on Dominica, Lesser Antilles. Chave J (Ed.). PLoS ONE 6: e18790. https://doi.org/10.1371/journal.pone.0018790

NUÑEZ MA, RELVA MA, SIMBERLOFF D (2008) Enemy release or invasional meltdown? Deer preference for exotic and native trees on Isla Victoria, Argentina. Austral Ecology 33: 317–323. https://doi.org/10.1111/j.1442-9993.2007.01819.x

Nuñez MA, Simberloff D, Relva MA (2008) Seed predation as a barrier to alien conifer invasions. Biological Invasions 10: 1389–1398. https://doi.org/10.1007/s10530-007-9214-x

Nylund GM, Pereyra RT, Wood HL, Johannesson K, Pavia H (2012) Increased resistance towards generalist herbivory in the new range of a habitat-forming seaweed. Ecosphere 3: art125. https://doi.org/10.1890/es12-00203.1

Oduor AMO, Kleunen M van, Stift M (2017) In the presence of specialist root and shoot herbivory, invasive-range Brassica nigra populations have stronger competitive effects than native-range populations. Buckley Y (Ed.). Journal of Ecology 105: 1679–1686. https://doi.org/10.1111/1365-2745.12779

Oduor AMO, Lankau RA, Strauss SY, Gómez JM (2011) Introduced Brassica nigra populations exhibit greater growth and herbivore resistance but less tolerance than native populations in the native range. New Phytologist 191: 536–544. https://doi.org/10.1111/j.1469-8137.2011.03685.x

PAN X-Y, JIA X, FU D-J, LI B (2013) Geographical diversification of growth–defense strategies in an invasive plant. Journal of Systematics and Evolution 51: 308–317. https://doi.org/10.1111/j.1759-6831.2012.00239.x

Park MG, Blossey B (2008) Importance of plant traits and herbivory for invasiveness of Phragmites australis (Poaceae). American Journal of Botany 95: 1557–1568. https://doi.org/10.3732/ajb.0800023

Parker IM, Gilbert GS (2007) WHEN THERE IS NO ESCAPE: THE EFFECTS OF NATURAL ENEMIES ON NATIVE, INVASIVE, AND NONINVASIVE PLANTS. Ecology 88: 1210–1224. https://doi.org/10.1890/06-1377

Parker JD, Hay ME (2005) Biotic resistance to plant invasions? Native herbivores prefer non-native plants. Ecology Letters 8: 959–967. https://doi.org/10.1111/j.1461-0248.2005.00799.x

Pearson DE, Callaway RM, Maron JL (2011) Biotic resistance via granivory: establishment by invasive, naturalized, and native asters reflects generalist preference. Ecology 92: 1748–1757. https://doi.org/10.1890/11-0164.1

Pedersen MF, Johnsen KL, Halle LL, Karling ND, Salo T (2016) Enemy release an unlikely explanation for the invasive potential of the brown alga Sargassum muticum: experimental results, literature review and meta-analysis. Marine Biology 163. https://doi.org/10.1007/s00227-016-2968-x

Pirk GI, Farji-Brener AG (2012) Foliar herbivory and its effects on plant growth in native and exotic species in the Patagonian steppe. Ecological Research 27: 903–912. https://doi.org/10.1007/s11284-012-0968-y

Porazinska DL, Pratt PD, Glblin-Davis RM (2007) Consequences of Melaleuca quinquenervia Invasion on Soil Nematodes in the Florida Everglades. Journal of Nematology 39: 305–312.

Prider J, Watling J, Facelli JM (2008) Impacts of a native parasitic plant on an introduced and a native host species: implications for the control of an invasive weed. Annals of Botany 103: 107–115. https://doi.org/10.1093/aob/mcn214

Prior KM, Hellmann JJ (2013) Does enemy loss cause release? A biogeographical comparison of parasitoid effects on an introduced insect. Ecology 94: 1015–1024. https://doi.org/10.1890/12-1710.1

Putten WH van der, Yeates GW, Duyts H, Reis CS, Karssen G (2005) Invasive plants and their escape from root herbivory: a worldwide comparison of the root-feeding nematode communities of the dune grass Ammophila arenaria in natural and introduced ranges. Biological Invasions 7: 733–746. https://doi.org/10.1007/s10530-004-1196-3

Putten WH van der, Kowalchuk GA, Brinkman EP, Doodeman GTA, Kaaij RM van der, Kamp AFD, Menting FBJ, Veenendaal EM (2007) SOIL FEEDBACK OF EXOTIC SAVANNA GRASS RELATES TO PATHOGEN ABSENCE AND MYCORRHIZAL SELECTIVITY. Ecology 88: 978–988. https://doi.org/10.1890/06-1051

Qi S-S, Liu Y-J, Dai Z-C, Wan L-Y, Du D-L, Ju R-T, Wan JSH, Bonser SP (2019) Allelopathy confers an invasive Wedelia higher resistance to generalist herbivore and pathogen enemies over its native congener. Oecologia 192: 415–423. https://doi.org/10.1007/s00442-019-04581-z

Radho-Toly S, Majer JD, Yates C (2001) Impact of fire on leaf nutrients, arthropod fauna and herbivory of native and exotic eucalypts in Kings Park, Perth, Western Australia. Austral Ecology 26: 500–506. https://doi.org/10.1046/j.1442-9993.2001.01133.x

Reddy AM, Carruthers RI, Mills NJ (2015) No evolution of reduced resistance and compensation for psyllid herbivory by the invasive Genista monspessulana. Plant Ecology 216: 1457–1468. https://doi.org/10.1007/s11258-015-0525-1

Reinhart KO, Callaway RM (2004) SOIL BIOTA FACILITATE EXOTIC ACER INVASIONS IN EUROPE AND NORTH AMERICA. Ecological Applications 14: 1737–1745. https://doi.org/10.1890/03-5204

Reinhart KO, Packer A, Putten WHV der, Clay K (2003) Plant-soil biota interactions and spatial distribution of black cherry in its native and invasive ranges. Ecology Letters 6: 1046–1050. https://doi.org/10.1046/j.1461-0248.2003.00539.x

Reinhart KO, Tytgat T, Putten WHV der, Clay K (2010) Virulence of soil-borne pathogens and invasion by Prunus serotina. New Phytologist 186: 484–495. https://doi.org/10.1111/j.1469-8137.2009.03159.x

Roche DG, Leung B, Franco EFM, Torchin ME (2010) Higher parasite richness, abundance and impact in native versus introduced cichlid fishes. International Journal for Parasitology 40: 1525–1530. https://doi.org/10.1016/j.ijpara.2010.05.007

Rode NO, Lievens EJP, Segard A, Flaven E, Jabbour-Zahab R, Lenormand T (2013) Cryptic microsporidian parasites differentially affect invasive and native Artemia spp. International Journal for Parasitology 43: 795–803. https://doi.org/10.1016/j.ijpara.2013.04.009

Rodríguez J, Novoa A, Cordero-Rivera A, Richardson DM, González L (2020) Biogeographical comparison of terrestrial invertebrates and trophic feeding guilds in the native and invasive ranges of Carpobrotus edulis. NeoBiota 56: 49–72. https://doi.org/10.3897/neobiota.56.49087

Roets F, Benadé PC, Samways MJ, Veldtman R (2018) Better colony performance, not natural enemy release, explains numerical dominance of the exotic Polistes dominula wasp over a native congener in South Africa. Biological Invasions 21: 925–933. https://doi.org/10.1007/s10530-018-1870-5

Roy BA, Coulson T, Blaser W, Policha T, Stewart JL, Blaisdell GK, Güsewell S (2011) Population regulation by enemies of the grass Brachypodium sylvaticum: demography in native and invaded ranges. Ecology 92: 665–675. https://doi.org/10.1890/09-2006.1

Sagerman J, Enge S, Pavia H, Wikström SA (2015) Low feeding preference of native herbivores for the successful non-native seaweed Heterosiphonia japonica. Marine Biology 162: 2471–2479. https://doi.org/10.1007/s00227-015-2730-9

Salonen JK, Marjomäki TJ, Taskinen J (2016) An alien fish threatens an endangered parasitic bivalve: the relationship between brook trout Salvelinus fontinalis and freshwater pearl mussel Margaritifera margaritifera in northern Europe. Aquatic Conservation: Marine and Freshwater Ecosystems 26: 1130–1144. https://doi.org/10.1002/aqc.2614

Sánchez MI, Rode NO, Flaven E, Redón S, Amat F, Vasileva GP, Lenormand T (2012) Differential susceptibility to parasites of invasive and native species of Artemia living in sympatry: consequences for the invasion of A. franciscana in the Mediterranean region. Biological Invasions 14: 1819–1829. https://doi.org/10.1007/s10530-012-0192-2

Sarabeev V (2015) Helminth species richness of introduced and native grey mullets (Teleostei: Mugilidae). Parasitology International 64: 6–17. https://doi.org/10.1016/j.parint.2015.01.001

Schierenbeck KA, Mack RN, Sharitz RR (1994) Effects of Herbivory on Growth and Biomass Allocation in Native and Introduced Species of Lonicera. Ecology 75: 1661–1672. https://doi.org/10.2307/1939626

Schoeman AL, Kruger N, Secondi J, Preez LH du (2019) Repeated reduction in parasite diversity in invasive populations of Xenopus laevis: a global experiment in enemy release. Biological Invasions 21: 1323–1338. https://doi.org/10.1007/s10530-018-1902-1

Schrieber K, Wolf S, Wypior C, Höhlig D, Keller SR, Hensen I, Lachmuth S (2019) Release from natural enemies mitigates inbreeding depression in native and invasive Silene latifolia populations. Ecology and Evolution 9: 3564–3576. https://doi.org/10.1002/ece3.4990

Schutzenhofer MR (2007) The Effect of Herbivory on the Mating System of Congeneric Native and Exotic Lespedeza Species. International Journal of Plant Sciences 168: 1021–1026. https://doi.org/10.1086/518941

Schutzenhofer MR, Valone TJ, Knight TM (2009) Herbivory and population dynamics of invasive and native Lespedeza. Oecologia 161: 57–66. https://doi.org/10.1007/s00442-009-1354-5

Schwartz N, Rohde S, Hiromori S, Schupp PJ (2016) Understanding the invasion success of Sargassum muticum: herbivore preferences for native and invasive Sargassum spp. Marine Biology 163. https://doi.org/10.1007/s00227-016-2953-4

Siemann E, Rogers WE (2003) Herbivory, Disease, Recruitment Limitation, and Success of Alien and Native Tree Species. Ecology 84: 1489–1505. https://doi.org/10.1890/0012-9658(2003)084[1489:HDRLAS]2.0.CO;2

Siemann E, Rogers WE (2006) Recruitment Limitation, Seedling Performance and Persistence of Exotic Tree Monocultures. Biological Invasions 8: 979–991. https://doi.org/10.1007/s10530-005-0825-9

Siemann E, Rogers WE, Dewalt SJ (2006) Rapid adaptation of insect herbivores to an invasive plant. Proceedings of the Royal Society B: Biological Sciences 273: 2763–2769. https://doi.org/10.1098/rspb.2006.3644

Siemann E, DeWalt SJ, Zou J, Rogers WE (2016) An experimental test of the EICA Hypothesis in multiple ranges: invasive populations outperform those from the native range independent of insect herbivore suppression. AoB Plants: plw087. https://doi.org/10.1093/aobpla/plw087

Simoncini M, Miller RJ (2007) Feeding preference of Strongylocentrotus droebachiensis (Echinoidea) for a dominant native ascidian, Aplidium glabrum, relative to the invasive ascidian Botrylloides violaceus. Journal of Experimental Marine Biology and Ecology 342: 93–98. https://doi.org/10.1016/j.jembe.2006.10.019

Southwood TRE, Moran VC, Kennedy CEJ (1982) The Richness, Abundance and Biomass of the Arthropod Communities on Trees. The Journal of Animal Ecology 51: 635. https://doi.org/10.2307/3988

STASTNY M, SCHAFFNER U, ELLE E (2005) Do vigour of introduced populations and escape from specialist herbivores contribute to invasiveness? Journal of Ecology 93: 27–37. https://doi.org/10.1111/j.1365-2745.2004.00962.x

Strauss SY, Stanton ML, Emery NC, Bradley CA, Carleton A, Dittrich-Reed DR, Ervin OA, Gray LN, Hamilton AM, Rogge JH, Harper SD, Law KC, Pham VQ, Putnam ME, Roth TM, Theil JH, Wells LM, Yoshizuka EM (2009) Cryptic seedling herbivory by nocturnal introduced generalists impacts survival, performance of native and exotic plants. Ecology 90: 419–429. https://doi.org/10.1890/07-1533.1

Stricker KB, Stiling P (2013) Release from herbivory does not confer invasion success for Eugenia uniflora in Florida. Oecologia 174: 817–826. https://doi.org/10.1007/s00442-013-2798-1

Stutz S, Štajerová K, Hinz HL, Müller-Schärer H, Schaffner U (2016) Can enemy release explain the invasion success of the diploid Leucanthemum vulgare in North America? Biological Invasions 18: 2077–2091. https://doi.org/10.1007/s10530-016-1152-z

Sullivan JJ, Winks CJ, Fowler SV (2008) Novel host associations and habitats for Senecio-specialist herbivorous insects in Auckland. New Zealand Journal of Ecology 32: 219–224.

Tewes LJ, Mueller C (2018) Syndromes in suites of correlated traits suggest multiple mechanisms facilitating invasion in a plant range-expander. NeoBiota 37: 1–22. https://doi.org/10.3897/neobiota.37.21470

Tomas F, Box A, Terrados J (2010) Effects of invasive seaweeds on feeding preference and performance of a keystone Mediterranean herbivore. Biological Invasions 13: 1559–1570. https://doi.org/10.1007/s10530-010-9913-6

Tuttle LJ, Sikkel PC, Cure K, Hixon MA (2016) Parasite-mediated enemy release and low biotic resistance may facilitate invasion of Atlantic coral reefs by Pacific red lionfish (Pterois volitans). Biological Invasions 19: 563–575. https://doi.org/10.1007/s10530-016-1342-8

Valverde PL, Arroyo J, Núñez-Farfán J, Castillo G, Calahorra A, Pérez-Barrales R, Tapia-López R (2015) Natural selection on plant resistance to herbivores in the native and introduced range. AoB Plants 7: plv090. https://doi.org/10.1093/aobpla/plv090

Vasquez EC, Meyer GA (2010) Relationships among leaf damage, natural enemy release, and abundance in exotic and native prairie plants. Biological Invasions 13: 621–633. https://doi.org/10.1007/s10530-010-9853-1

Vermeij MJA, Smith TB, Dailer ML, Smith CM (2008) Release from native herbivores facilitates the persistence of invasive marine algae: a biogeographical comparison of the relative contribution of nutrients and herbivory to invasion success. Biological Invasions 11: 1463–1474. https://doi.org/10.1007/s10530-008-9354-7

Veselkin DV, Kuyantseva NB, Chashchina OE, Mumber AG, Zamshina GA, Molchanova DA (2019) Levels of Leaf Damage by Phyllophages in Invasive Acer negundo and Native Betula pendula and Salix caprea. Russian Journal of Ecology 50: 511–516. https://doi.org/10.1134/s1067413619060134

Vignon M, Sasal P, Galzin R (2008) Host introduction and parasites: a case study on the parasite community of the peacock grouper Cephalopholis argus (Serranidae) in the Hawaiian Islands. Parasitology Research 104: 775–782. https://doi.org/10.1007/s00436-008-1254-3

Vilà M, Maron JL, Marco L (2004) Evidence for the enemy release hypothesis in Hypericum perforatum. Oecologia 142: 474–479. https://doi.org/10.1007/s00442-004-1731-z

Volin JC, Kruger EL, Volin VC, Tobin MF, Kitajima K (2009) Does release from natural belowground enemies help explain the invasiveness of Lygodium microphyllum? A cross-continental comparison. Plant Ecology 208: 223–234. https://doi.org/10.1007/s11258-009-9700-6

Wan JSH, Bonser SP (2016) Enemy release at range edges: do invasive species escape their herbivores as they expand into new areas? Journal of Plant Ecology 9: 636–647. https://doi.org/10.1093/jpe/rtw003

Wandrag EM, Sheppard AW, Duncan RP, Hulme PE (2014) Pollinators and predators at home and away: do they determine invasion success for Australian Acacia in New Zealand? Kissling WD (Ed.). Journal of Biogeography 42: 619–629. https://doi.org/10.1111/jbi.12455

Wang R-F, Feng Y-L (2016) Tolerance and resistance of invasive and native Eupatorium species to generalist herbivore insects. Acta Oecologica 77: 59–66. https://doi.org/10.1016/j.actao.2016.09.001

Wang S, Wang G, Weinberger F, Bian D, Nakaoka M, Lenz M (2016) Anti-epiphyte defences in the red seaweed Gracilaria vermiculophylla: non-native algae are better defended than their native conspecifics. Gibson D (Ed.). Journal of Ecology 105: 445–457. https://doi.org/10.1111/1365-2745.12694

Wang Y-J, Chen D, Yan R, Yu F-H, Kleunen M van (2019) Invasive alien clonal plants are competitively superior over co-occurring native clonal plants. Perspectives in Plant Ecology, Evolution and Systematics 40: 125484. https://doi.org/10.1016/j.ppees.2019.125484

Wikström SA, Steinarsdóttir MB, Kautsky L, Pavia H (2006) Increased chemical resistance explains low herbivore colonization of introduced seaweed. Oecologia 148: 593–601. https://doi.org/10.1007/s00442-006-0407-2

Williams JL, Auge H, Maron JL (2010) Testing hypotheses for exotic plant success: parallel experiments in the native and introduced ranges. Ecology 91: 1355–1366. https://doi.org/10.1890/08-2142.1

Williams V-RJ, Sahli HF (2016) A Comparison of Herbivore Damage on Three Invasive Plants and Their Native Congeners: Implications for the Enemy Release Hypothesis. Castanea 81: 128–137. https://doi.org/10.2179/15-069

Wolfe LM (2002) Why Alien Invaders Succeed: Support for the Escape-from-Enemy Hypothesis. The American Naturalist 160: 705–711. https://doi.org/10.1086/343872

Wolfe LM, Elzinga JA, Biere A (2004) Increased susceptibility to enemies following introduction in the invasive plant Silene latifolia. Ecology Letters 7: 813–820. https://doi.org/10.1111/j.1461-0248.2004.00649.x

Wróbel A, Crone EE, Zwolak R (2019) Differential impacts of soil microbes on native and co-occurring invasive tree species. Ecosphere 10. https://doi.org/10.1002/ecs2.2802

XIONG W, YU D, WANG Q, LIU C, WANG L (2008) A snail prefers native over exotic freshwater plants: implications for the enemy release hypotheses. Freshwater Biology 53: 2256–2263. https://doi.org/10.1111/j.1365-2427.2008.02058.x

Yan YZ, Chen YF (2007) Changes in the life history of Abbottina rivularis in Lake Fuxian. Journal of Fish Biology 70: 959–964. https://doi.org/10.1111/j.1095-8649.2007.01343.x

Yun HY, Molis M (2012) Comparing the ability of a non-indigenous and a native seaweed to induce anti-herbivory defenses. Marine Biology 159: 1475–1484. https://doi.org/10.1007/s00227-012-1926-5

Zas R, Moreira X, Sampedro L (2011) Tolerance and induced resistance in a native and an exotic pine species: relevant traits for invasion ecology. Journal of Ecology 99: 1316–1326. https://doi.org/10.1111/j.1365-2745.2011.01872.x

Zhao Y-Z, Liu M-C, Feng Y-L, Wang D, Feng W-W, Clay K, Durden LA, Lu X-R, Wang S, Wei X-L, Kong D-L (2020) Release from below- and aboveground natural enemies contributes to invasion success of a temperate invader. Plant and Soil 452: 19–28. https://doi.org/10.1007/s11104-020-04520-5

Zheng Y-L, Feng Y-L, Wang R-F, Shi X-D, Lei Y-B, Han L-H (2012) Invasive Eupatorium adenophorum suffers lower enemy impact on carbon assimilation than native congeners. Ecological Research 27: 867–872. https://doi.org/10.1007/s11284-012-0964-2

Zocca A, Zanini C, Aimi A, Frigimelica G, Porta NL, Battisti A (2008) Spread of plant pathogens and insect vectors at the northern range margin of cypress in Italy. Acta Oecologica 33: 307–313. https://doi.org/10.1016/j.actao.2008.01.004

Zou J, Rogers WE, Siemann E (2007) Increased competitive ability and herbivory tolerance in the invasive plant Sapium sebiferum. Biological Invasions 10: 291–302. https://doi.org/10.1007/s10530-007-9130-0

Zou J, Siemann E, Rogers WE, DeWalt SJ (2008) Decreased resistance and increased tolerance to native herbivores of the invasive plant Sapium sebiferum. Ecography 31: 663–671. <https://doi.org/10.1111/j.0906-7590.2008.05540.x>

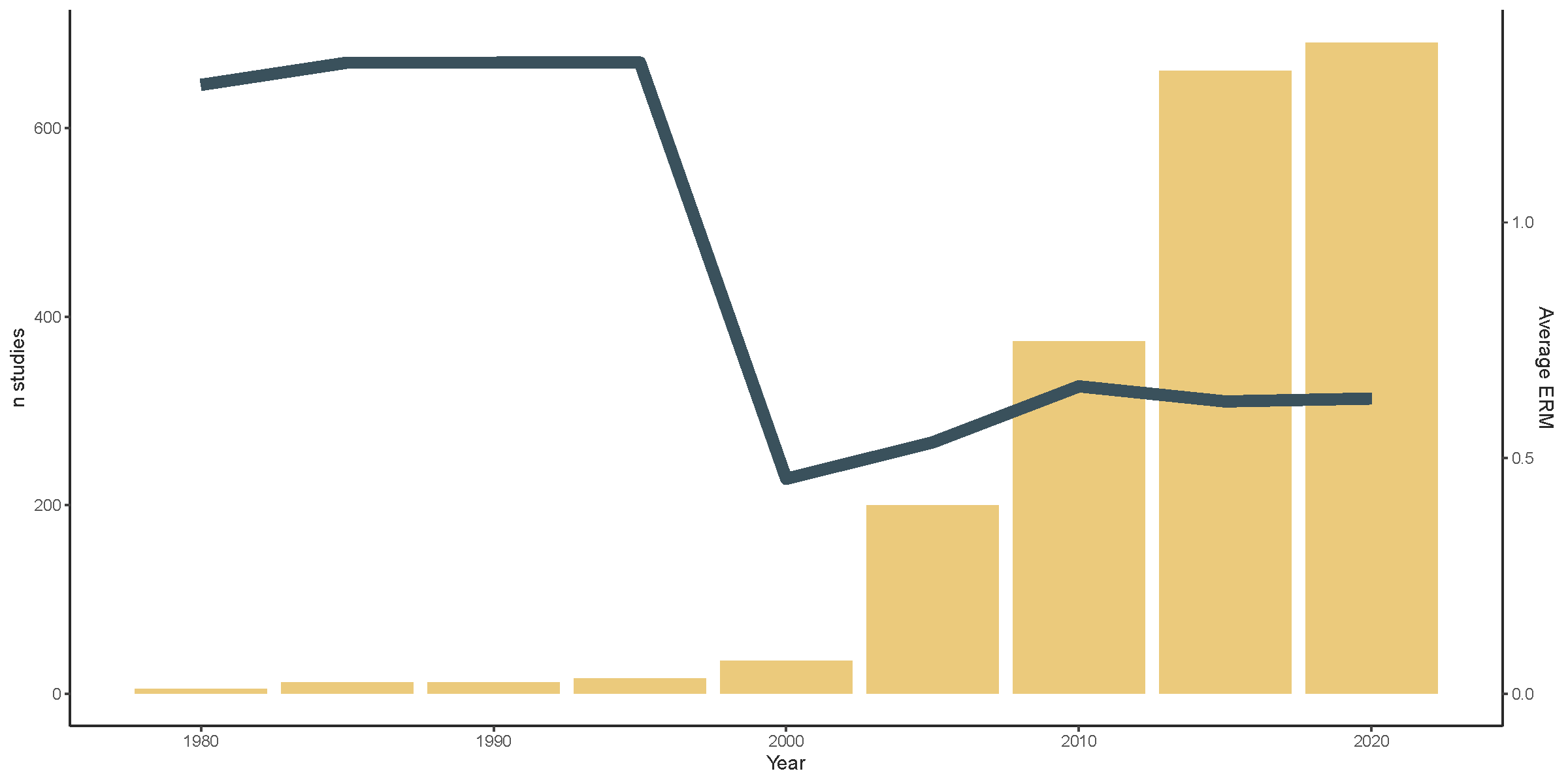

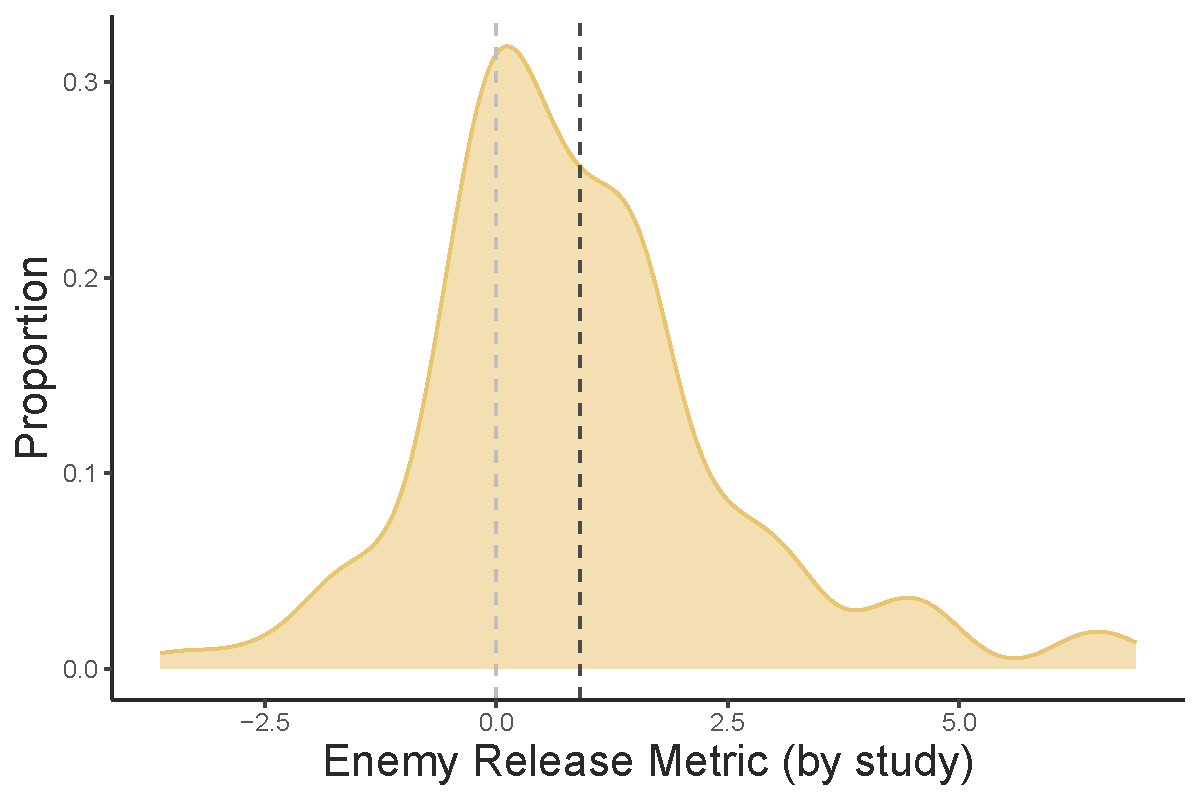

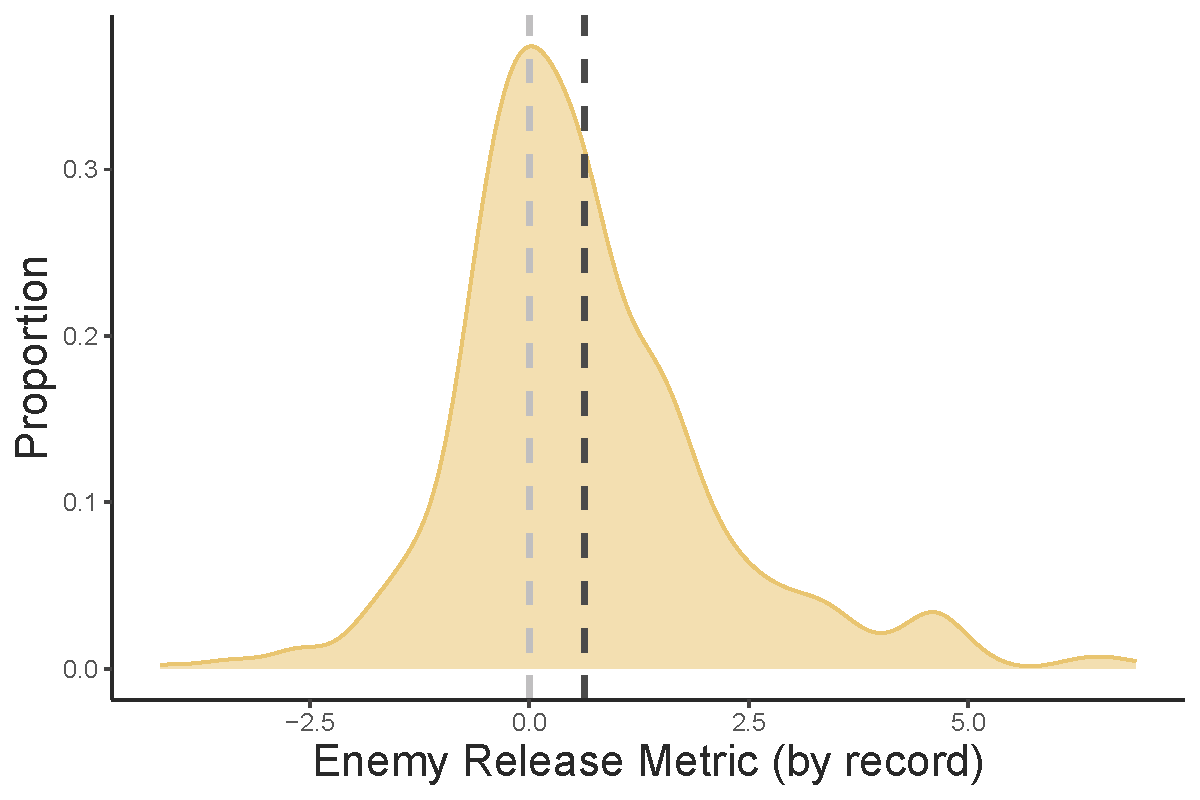

a)

b)

c)

**Figure 1.** Support for the enemy release hypothesis a) by individual record, b) by study and c) cumulative average support throughout time (line) as the number of enemy release studies accumulate (yellow bars). Lighter dotted line indicates zero whereas darker dotted line indicates mean.

The proportion of studies which found positive enemy release was higher than the proportion of population records with positive enemy release, as negative values are often overlooked.
